# A comparison of several media types and basic techniques used to assess outdoor airborne fungi in Melbourne, Australia

**DOI:** 10.1101/2020.08.27.269704

**Authors:** Wesley D Black

## Abstract

Despite the recent increase in interest in indoor air quality regarding mould, there is no single widely accepted standard media for the detection of airborne fungi, nor verification of many commonly used techniques. Commonly used media including malt-extract agar (MEA), Sabouraud dextrose agar (Sab, SDA, SabCG), potato dextrose agar (PDA) with and without antibiotics chloramphenicol & gentamycin (CG) were compared for their suitability in detecting a range of common airborne fungi by collecting 150 L outdoor air on a number of different days and seasons via an Anderson 400-hole sampler in suburban Melbourne, Australia. There was relatively little variation in mean numbers of colony forming units (CFU) and types of fungi recovered between MEA, PDA, SabCG media groups relative to variation within each group. There was a significant difference between SabCG, Dichloran-18% glycerol (DG18) and V8® Original juice agar media, however. Antibiotics reliably prevented the growth of bacteria that typically interfered with the growth and appearance of fungal colonies. There was no significant evidence for a growth enhancing factor from potato, mineral supplements or various vegetable juices. Differing glucose concentrations had modest effects, showing a vague ideal at 2%-4% with peptone. Sanitisation/sterilisation of the aluminium Andersen 400-hole sampler top-plate by flame is possible, but not strictly required nor advisable. The use of SabCG as a standard medium was generally supported.

## Introduction

Mould are the wide range of fungal organisms that flourish under damp conditions indoors and outdoors, and in humans exposure is linked to the exacerbation of asthma, allergic rhinitis and occasionally infection. Intoxication from the ingestion of mould/mycotoxin-contaminated foods is known [1], but the causal relationship between mould inhalation and noted significant respiratory conditions including acute infant idiopathic pulmonary haemorrhage requires additional investigation [2].

A review of the current literature suggests there is no universally or even widely-accepted method for detecting, identifying and/or enumerating them within buildings, and similarly a lack of widely-accepted limits for maximum permissible and/or normal exposure to occupants, or even what may constitute a ‘mouldy’ house.

Outdoors, various moulds, yeasts, various other fungi and organisms saprophytically degrade organic matter such as fallen leaves, trees, etc., and are generally ecologically beneficial [2]. The most common outdoor mould & yeast genera/types noted in studies in the Northern Hemisphere were *Cladosporium, Aspergillus, Penicillium, Alternaria, Candida, Botrytis and Helminthosporium* [2]. Within houses not known to be problematic the most common mould & yeast genera/types noted were very similar to outdoors, but included *Epicoccum* mould and *Streptomyces* bacteria [2]. These indoor organisms are not usually a problem except in persistently humid or wet areas of houses in which such organisms significantly grow in number [2]. Exposures to mould varies depending on a range of factors including regional differences, local climate including outdoor humidity and wind, shade, organic debris, landscape maintenance, etc., heating and cooling systems, indoor humidity and air-filtration and ventilation systems [2]. Dampness in a house is also associated with the deterioration of structural components such as plasterboard / Gyprock / drywall panels [3].

Mould has the potential to cause a variety of adverse health effects by both immune- and non-immune-related mechanisms including immunoglobulin E mediated responses and allergic rhinitis, conjunctivitis and asthma, allergic bronchopulmonary aspergillosis, allergic fungal sinusitis and hypersensitivity pneumonitis, and actual infection, respectively [2]. Other fungi including yeasts often found in houses and foodstuffs have been noted as emerging pathogens [4], including *Rhodotorula*, often seen as the pink stain on tile grout in bathroom showers.

There are established health risks associated with living in damp indoor environments per se, including respiratory symptoms such as wheezing, coughing and allergic rhinitis / ‘hay fever,’ and asthma symptoms in sensitised persons [2]. It is likely the dampness leads to excess fungal growth indoors and consequent exposure of occupants, but the precise mechanism remains unclear as yet.

While fungi are ubiquitous outdoors and impractical to prevent from being blown into a normal house, it appears the challenge is to prevent them from actively colonising and growing within the house, and the key to this is to control indoor moisture by various means [2]. Once a building is wet enough for such colonisation, remediation is required promptly to prevent further growth by way of thorough drying ideally within 24-72 hours. Failing this, thorough physical removal of the then numerous fungal particles including spores and non-spore fragments is also required to reduce exposure to occupants and site workers [5,6]. There are significant differences of opinion globally on how best to achieve this, and to what degree, and how to objectively determine if it has actually been achieved [2,7–13]. This is curious given the increasing number of legal disputes at least in the state of Victoria, Australia [14] and the Australian Federal Government interest [15].

The main established means of determining if a building has been water-damaged and mouldy is to compare air samples taken from outdoors and in a number of locations indoors, counting fungal particles by either culture-based methods for ‘viable’ colony-forming units (CFU), or microscopy-based ‘total-count’ of identifiable fungal particles, or ideally both in order to overcome the limitations of each, as also airborne and settled particles, and measurement of excess dampness and humidity [7,16].

A long-used method of collecting and enumerating airborne viable particles is the Anderson air sampler [17]. The unit consists essentially of a calibrated air-pump drawing air through an airtight disk-like top-plate assembly with a set of 200 or 400 small holes directly over a 90 mm diameter Petri dish of suitable agar gel media held within the assembly. Air drawn through the holes hits the gel surface, deflecting sharply and hence depositing airborne particles on the damp gel. The Petri dish is then removed from the assembly and incubated for a period to allow growth of organisms able to do so on that media at the incubation temperature, and facilitating identification and enumeration.

The collection and enumeration of surface-borne or settled viable particles is often by simple sterile swab collection of a known surface area, then transferred to a similar Petri dish of media and incubated. Other methods exist such as replicate organism detection and counting (RODAC) touch-plates that employ a slightly convex agar gel surface with flattened top that is applied to the test surface then incubated [18,19]. These, however, have been found to have a poor and variable recovery rate from standardised indicator-organism seeded surfaces between several commercial products [20]. It also remains to be seen which agar media is best suited to the purpose of viable-counts in general regardless of the method of sample collection.

The use of settle-plates or open Petri dishes exposed to air for a time to enumerate viable airborne/settling particles has been used in the past [21] or more recently by researchers without ready access to powered air-sampling devices [22–25].

Methods of estimating the degree of microbial contamination of surfaces by the detection of adenosine triphosphate (ATP), the ‘universal energy currency’ of a living organism is established for industries such as food preparation surfaces and machinery, surgical operating theatres and sterile containment facilities for the manufacture of pharmaceuticals [26]. Such methods, however, were developed more for the detection of actively respiring easily lyseable organisms such as vegetative bacterial cells and yeasts on surfaces potentially contaminated with foodstuffs more as a means of determining the efficacy of cleaning and sanitisation protocols. Such systems have been criticised, however for not considering the significant variation in ATP content between the range of organisms on any given surface, and their states of nutrition, growth cycle, sporulation/germination, and the significant relative amount of ATP from fresh foodstuff residues themselves on the same surface [26]. More recently, a test that detects ATP, and adenosine diphosphate (ADP) and adenosine monophosphate (AMP) was developed and is purported to be more sensitive and reliable [27].

Airborne particles can be detected and enumerated by other means including LASER particle counters that monitor the number of deflections and angle or amount of deflection (and hence the size of the particle) from particles passing through the measurement chamber as they are drawn through it [28,29]. While widely used in various industrial applications to monitor dusts, such devices are unable to determine if a particle is a viable spore, nonviable spore, other whole organisms, pollen, mineral grit, sawdust, skin flakes / dander, hair, animal fur, feathers, jumper/sweater fluff, etc.

A pilot investigation conducted in the subtropical-oceanic city, Brisbane, Australia, found no statistically significant associations between fungal spore concentrations and sub-micrometre particle concentrations [30]. This is of considerable practical nuisance in a normal inhabited house compared to perhaps an industrial clean-room or sterile pharmaceutical dispensing suite in which any such particles are detrimental and kept below prescribed limits such as the ISO 14644-1 Cleanroom Standards [31].

Regardless, the detection of viable fungi by culture for enumeration and identification seems an important aspect of assessing a damp building that other methods are unable to quite address. Other studies have described ‘indicator’ moulds to attempt to estimate how mouldy a house is [32]. This is an extension of a long-established concept in microbiology in which an ‘indicator organism’ is used to determine originally if a sample is positive for faeces in which many pathogens may not be present, and are often difficult to detect even when present, and thus indicator organisms are used instead that are known to virtually always be present in faeces and are more easily and reliably detected, but may not actually be pathogenic themselves. In this case the indicator moulds were used to determine that there was a correlation between noted dampness, visible signs of mould and damage, and detectable mouldiness [32]. A study of dust extracted from carpets and rugs in many houses in Wallaceburg, Ontario, Canada, found that *Alternaria alternaria, Aureobasidium pullulans, Eurotium herbariorum, Epicoccum nigrum, Aspergillus versicolor* and *Penicillium chrysogenum* were present in 50% or more of the samples analysed using Rose Bengal agar medium (RBA) with antibiotics, with and without 25% glycerol, however [33].

Various studies have used various media for various purposes. A study of 80 not-notably-mouldy living areas and 14 notably-mouldy rooms in Germany used dichloran-18%-glycerol agar (DG18) and malt-extract agar (MEA) media [32]. While the study did draw a correlation, it remains to be seen that these are the ideal media for these types of studies given they appear to be used more by tradition than their demonstrated suitability for use in a variety of conditions, locations and climates, or the specific purpose of assessing indoor air quality. DG18 is used as a selective media for xerotolerant and mesophilic organisms given its low water-activity (a_w_) courtesy of its high salt/solute content [34].

MEA media have been used for some 100 years at least, presumably because of its relative low cost and the high availability of malt extract, and presumably the likely common utilisation of the dominant sugar, maltose, by organisms rotting or fermenting grains. This was hence a likely subject of interest to early microbiologists, farmers, bakers and brewers. It is also known that some brewing / baking yeast (*Saccharomyces cerevisiae*) strains have various utilisation of maltose [35]. It however remains to be seen that the range of various other fungi growing in other micro-environments and ecological niches including skin, compost, rotting leaf-litter and damp cardboard or carpets in water damaged buildings (WDB) would also utilise maltose given its likely typical absence.

A study of 64 homes in the UK used Sabouraud 4% glucose chloramphenicol agar media (SabC, or Sabouraud dextrose agar with chloramphenicol, SDAC) via an Andersen 6-stage sampler, and did establish a correlation between visible mould and detected mould, albeit with self-noted wide variance, and noted concerns about the variability of indoor air velocity and activities acting to suspend dusts and hence increase data uncertainty [36].

Sabouraud media was originally formulated well before the discovery of antibiotics for the cultivation of dermophyte fungi associated with skin, nail, oral, respiratory and urogenital conditions. This required a medium able to reliably grow a range of fungi but ideally not the vast number of otherwise harmless resident bacteria, and without the use of antibiotics given the era of development. Given that a large range of bacteria do not grow well in somewhat acid media but relatively many fungi do, the original formulation employed an acidic pH plus the fairly universal energy source, glucose, and a general amino acid / peptide supplement, peptone, made from a digest of proteins from various sources, such as Mycological Peptone Powder (Oxoid, LP0040), among others. A study comparing variations of Sabouraud recipes for the isolation of fungi from the sputum of patients with cystic fibrosis found that a slightly lesser amount of glucose (16.7 g/L) plus yeast extract (30 g/L) and peptone (6.8 g/L) adjusted to pH 6.3 and including a range of antibiotics was the most sensitive medium tested for that specific application [37], but remains to be seen regarding indoor/outdoor airborne fungi.

Other media commonly used includes DG18, with dichloran added to limit the spread of some fast-growing fungal colonies, limiting their diameter [38] and reducing the problem of covering over other, smaller colonies obfuscating them and making identification and enumeration difficult. DG18 has a relatively low a_w_ via the inclusion of salts and 18% glycerol [34,39]. The dichloran appears however to affect the growth of various fungi differently, barely limiting the growth of some while completely inhibiting others [39] and is somewhat toxic and hence somewhat less than ideal for handling and disposal [40].

Studies comparing various media used to sample indoor air fungi found that DG18 at 25°C generally recovered significantly higher numbers c.f., MEA and/or at 37°C [41]. Other workers noted that *Cladosporium halotolerans* more often survived sudden rehydration on high a_w_ media after having been dried and cultured in low a_w_ media than did *Aspergillus niger* and *Penicillium rubens* given these tended essentially to explode on rehydration [42]. Others suggested that the lower a_w_ of DG18 (∼0.96) c.f., other common media (∼0.99) may allow better recovery of food spoilage yeasts that originally grew at low a_w_, presumably having high internal osmolarity and hence at risk of explosion on high a_w_ media [39]. For context, typical seawater has an a_w_ of 0.98 [43], and many sea salt-preserved foods have an a_w_ of around 0.95 [44].

Studies of dust-borne fungi in houses using media including DG18 suggested that spores may be unable to grow in culture for various reasons including inappropriate nutrients, temperature or inhibitors, and hence yielding only loose associations in numbers of mould detected in notably-mouldy and non-mouldy houses [45,46]. Given that DG18 was originally developed to enumerate xerophilic foodborne moulds and yeasts this is unsurprising [34] and it was also found that it was less able to reliably culture various food spoilage fungi compared to other newer media and hence is not now recommended even for this purpose [39].

Rose Bengal medium is also used [38,39], being both a selective and differential medium in that the Rose Bengal dye is taken up by fungi more than other organisms and hence becoming pink/red in colour, but has the notable problem of the dye becoming toxic when exposed to light [47] and hence likely introducing greater variability in results. This is not an uncommon effect of dyes [48]. This medium appears useful more for the selection of mesophilic or xerophilic/xerotolerant fungi in a sample rather than the enumeration/estimation of the entire range of fungi including those requiring high a_w_, such as those found in a recently water-damaged building (WDB). There is evidence of a shift in fungal ecology between the outdoor environment, not visibly mouldy dwellings and visibly mouldy dwellings, becoming less relatively diverse presumably because of the overgrowth of fungi especially suited to the dwelling’s exact dampness, humidity, temperature, etc. [49,50].

Some moulds found in WDB such as *Stachybotrys chartarum atra* and *Chaetomium globosum* require very high a_w_, nearing total saturation, and may substantially lose their viability soon after collection from their active-growth site, and are relatively very slow growing, very often being totally obscured by faster-growing ‘early coloniser’ mould like *Penicillium, Aspergillus, Ulocladium*, etc., that that tend to spread over them [50]. Thus, a low a_w_ media may fail to detect these important indoor fungi.

A range of other media are also occasionally used for fungi for various objectives, including the passaging/sub-culturing of reference strains that have already been isolated and purified, and especially plant-borne pathogens (potato-dextrose agar, PDA) [51], food-spoilage fungi (DRBC, PCAC, TGYC, DG18) [39] or for selection/identification of fungi and bacteria able to utilise sucrose and inorganic nitrogen (Czapek-Dox Agar) [52] or enriching for specific subsets of microorganism populations (tap-water cellulose agar) [53], etc., which are hence significantly different from the objective of best estimating the numbers of viable fungi associated with WDB and normal indoor/outdoor airborne fungi.

The use of variously enriched media has been explored and does have some popularity among mycologists, including the addition of minerals and/or vegetable juices such as the commercially available ‘V8^®^ Original’ juice by the Campbell Soup Company, a blend of eight vegetables including mainly tomato juice, with the notion that there is some factor that enhances the growth and detection of fungi, albeit plant pathogens [32,50,54,55]. Some supplements such as molasses, V8^®^ juice, coconut, urea and ammonium variously affected the production of conidia, sclerotia and aflatoxins by *Aspergillus flavus* CA43 [56]. Other workers reported that the numbers of fungi recovered from houses varied over time when using media including Rose Bengal, MEA, V8 and DG18 agar that had an at least approximately 20% coefficient of variation [57].

The inclusion of anti-bacterial antibiotics in selective media for culturing fungi is now common depending on the application as bacteria and fungi compete for the same resources for their growth and hence affect the growth of other colonies nearby by either using limited resources faster and more effectively thus leaving little for competitors, or actively secreting substances that inhibit their growth, of which the antibiotic drug penicillin is a notable example, secreted by some strains/types of *Penicillium chrysogenum* mould in particular [58,59].

Some bacteria are known to secrete antifungal and/or antibacterial antibiotic compounds including chloramphenicol (chloromycetin), secreted by *Streptomyces venezuelae*, a Gram-positive soil bacterium [60]. Other notable findings were made during the development of antibacterial drugs [61].

Various *Lactobacillus* bacteria species are also known to have an antifungal effect against a range of fungi associated with vaginitis, onychomycosis and/or food spoilage [62,63]. *Bacillus subtilis* bacteria strains and other *Bacilli* have also been found with antifungal activity [64,65]. Several commercial preparations of probiotic capsules promoted as of benefit to people with various gastrointestinal complaints such as irritable bowel syndrome (IBS), abdominal pain/discomfort, flatulence/bloating, include a significant number of live/viable bacteria species noted as having antifungal activities such as *L. plantarum, L. casei, L. rhamnosus* and others [66].

Given that bacteria exist in vast numbers in ordinary topsoil, dusts, on skin, dander and hence the normal indoor environment, and they can affect the growth of fungi, antibiotics are often added to media used to analyse environmental samples. This is of less concern for media used to analyse typically sterile samples such as body tissues, blood and cerebrospinal fluid. Their relatively recent development has also meant that old media formulations did not include them, being either very expensive, not discovered yet, or often simply not stable enough to be autoclave sterilised at 121°C or stored in aqueous gel solution for a practical time period, or both, and penicillin is an example of this [58] as also are other labile antibiotics [61] in contrast to the bacterially produced anti-bacterial antibiotic chloramphenicol that is far more stable and hence suitable for such use [60].

It is also the case that while some bacteria can cause severe infections and presumably allergies when present in number in perhaps a formerly damp house, they are yet to be demonstrated as being airborne in quite the same manner and number that moulds and some other fungi are, which are usually well-adapted to this mode of dispersal [67–70].

## 2. Materials and methods

### Sabouraud / SabCG medium agar

as indicated, either pre-made complete Sabouraud Dextrose Agar powder (Oxoid CM0041, 65 g/L), mixed into cold reverse-osmosis (RO) purified water, adjusted to pH 5.6 ±0.2, autoclaved (121°C, 15 min, jacket off, no vacuum pre- or post-autoclave), cooled before adding 100x chloramphenicol-gentamycin stock (100xCG: 5 mg/mL chloramphenicol (Sigma C0378) in 50% ethanol, 40 mg/mL gentamicin (Pfizer / DBL / Pharmacia), hence 10 mL/L) before pouring. Otherwise where indicated, made ‘from scratch’ from individual components, being mycological peptone powder (Oxoid LP0040, 10 g/L), glucose powder (APS/AJAX, 40 g/L), bacteriological agar powder (Oxoid LP0011, 15 g/L) mixed into cold RO water to 1 L, adjusted to pH 5.6 ±0.2, autoclaved, cooled, with or without 100xGC addition as indicated then poured.

### PDA potato dextrose agar

Pre-made complete PDA powder (Oxoid CM0139, 39 g/L) mixed into cold RO water to 1 L, pH adjusted, autoclaved, cooled, with or without addition of 10 mL/L of 100xCG as indicated then poured.

### V8 media agar with chloramphenicol and gentamycin (V8c)

V8^®^ Original vegetable juice (Campbell Soup Company, Campbell’s Soup Australia, Lemnos, Victoria, 200 mL), calcium carbonate (Sigma, 2 g), bacteriological agar powder (15 g), mixed into cold RO water to 1 L, autoclaved, cooled, addition of 10 mL/L of 100xCG then poured.

### Dichloran 18% glycerol media agar with chloramphenicol and gentamycin (DG18c)

Pre-made dichloran glycerol agar base powder (Oxoid CM0729, 31.5 g), glycerol (Sigma, 176 mL) mixed into cold RO water to 1 L, autoclaved, cooled, addition of 10 mL / L of 100xCG added then poured.

### Malt extract media agar without/with chloramphenicol and gentamycin (MEA, MEACG)

malt-extract powder (Oxoid LP0039, 34 g/L), agar (10 g/L), mixed into cold tap water, adjusted to pH 5.5 ±0.2, autoclaved (110°C, 25 min), cooled, with or without 10 mL / L of 100xCG as indicated then poured.

### Peptone media agar with/without chloramphenicol and gentamycin (PeptoneCG, Peptone-only)

Mycological peptone powder (Oxoid LP0040, pH 5.3 at 2%), powdered agar and other components as indicated, cold RO water, autoclaved, cooled, with or without 10 mL / L of 100xCG as indicated and then poured.

### Maltose agar media

maltose monohydrate powder (Sigma M2250, 40 g/L) was added to liquid medias as indicated prior to being autoclaved, cooled, with or without 10 mL of 100xGC addition as indicated and then poured.

### Mineral supplement #1 (MS1) media agar

50⨯MS1 stock was prepared as 25 g CaCl_2_.2H_2_O (Sigma), RO water to 100 mL, then autoclave sterilised. MS1 medium was prepared as follows: 40 g glucose, 10 g peptone, 15 g bacteriological agar, RO water to 1 L. Autoclaved, cooled to approx. 50°C, 20 mL of 50xMS1 stock added dropwise with stirring of the liquid medium, 10 mL of 100xCG stock, pH adjusted to 6.7 +/- 0.3 and then poured.

### Mineral supplement #2 (MS2) media agar

50⨯MS2 stock was prepared as 40 g Ammonium dihydrogen phosphate (NH_4_)H_2_PO_4_, 5 g Potassium Chloride KCl, 5 g Magnesium Sulphate MgSO_4_.7H_2_O, 0.1 g Ferrous Sulphate FeSO_4_.7H_2_O, 0.1 g Zinc Sulphate ZnSO_4_.7H_2_O, 0.032 g Cupric Sulphate (anhydrous) CuSO_4_, RO Water to 100 mL then filter sterilised (Thermo Scientific™ 597-4520, 0.20 μm pore diameter). MS2 medium was prepared as per MS1 medium, but using the 50xMS2 stock.

### Vegetable supplements

Tomatoes (hydroponic ‘truss’ variety) and celery were bought fresh from a local supermarket (Sim’s IGA Supermarket, Footscray, Victoria, Australia) and each were juiced via kitchen food processor (Sunbeam). V8^®^ Original vegetable juice (Campbell Soup Company, in UHT sterilised bottles) were similarly acquired, being a combination of tomato, beets/beetroot, celery, carrot, lettuce, parsley, watercress, spinach juice concentrates and water.

### Clarification of vegetable juices

coarse filtration through clean/washed calico cloth, warming filtrate to approx. 50°C in a microwave oven, adding 1/4 volume of liquid agar stock (2% in RO water) that had been molten and cooled to approx. 50°C prior to addition, mixed then cooled to 4°C and filtered through clean calico cloth using some manual pressure, settled at 4°C for 30-60 minutes in a tall jar, decanted and filter sterilised (Corning 431218, 0.20 μm pore diameter).

### Heat-treatment of tomato juice

30 mL of freshly clarified sterile tomato juice in a sterile 50 mL plastic tube with a slightly loose lid was placed in approx. 200 mL boiling water in a borosilicate glass vessel and placed in a microwave oven, gently boiling for approx. 10 minutes, then cooled. Then 1 mL of each sterile supplement was added to the top of previously poured, set, room-temperature and slightly dried SabCG media plates and a sterile glass L-shaped bacteriology cell-spreader (Merck, S4522) was used to spread the liquid evenly over the entire surface and allowed to soak/dry before use.

### Air sampling

QuickTake30 (SKC Biosystems) with ‘SKC Biostage-400’ single-stage 400-hole sampler assembly attachment as per Andersen, 1958 [17] without any attachments atop the sampler top-plate, and at 1.5 m height, approx. 2 m from buildings, with wind-speeds approx. 2-5 knots on each occasion (i.e., still air and windy days were avoided, as also rain). Typically 150 L air samples were taken in triplicate in a ‘collated’ sequence, being all of the first plates of each different media, then all of the second plates, then all of the third plates, to best minimise the effect of differing air velocities and wind direction shifts, local dust-raising activities, etc.

### Bacteria powder

‘Double Strength Probiotic’ powder in capsules (Life-space Probiotics company), containing a combination of *Lactobacillus rhamnosus* Lr-32 & GG & HN001, *Bifidobacterium lactis* BI-4, *L. plantarum* Lp-115, *Streptococcus thermophilus* St-21, *L. casei* Lc-11, *L. paracasei* Lpc-37, *B. animalis* ssp. *lactis* HN019, *B. breve* Bb-03, *B. longum* Bl-05, *L. gasseri* Lg-36, *B. infantis* Bi-26, *L. delbrueckii* ssp. *bulgaricus* Lb-87 and *L. reuteri* 1E1, in approximately that order by number. A single capsule of supposedly 64 billion CFU/capsule was opened and a fine stream of the powder was blown by small electric fan towards the QuickTake30 sampling unit while in operation approximately 1 m away. This was intended only as an excess of bacteria known to interfere with fungal growth at >17,520 CFU/m^3^ air and hence at least one CFU for each of the 400 holes of the Andersen 400-hole sampler.

### CFU/m^3^

The term ‘airborne CFU/m^3^’ is used to generally describe viable airborne mould (*Actinomycetes, Zygomycetes*), yeasts, presumably also other fungi possibly including mushrooms, toadstools, earth-stars, puffballs, timber brown-rot, white-rot, etc., (*Basidiomycetes* generally), fungal plant pathogens and other organisms unaffected by chloramphenicol / gentamycin, slime-moulds and antibiotic-resistant bacteria able to grow at 27°C to a size detectable after 3 days on the stated media.

### Lugol’s Iodine

50 mg/mL iodine in 100 mg/mL potassium iodide solution as supplied in a standard Gram staining kit (Magnacol Pty Ltd, UK).

### Data processing and graphing

MS-Excel for Mac v16.35 (Microsoft). Analysis of variance (ANOVA) was by “Anova: Single Factor” [sic] method via Excel’s analysis tools package, as also mean and standard deviation (SD).

## Results

### Media tests: glucose, maltose, peptone and SabCG

Testing various malt-extract and maltose-based media (Fig 1) in late-January 2020 (summer) in Melbourne, Australia, indicated a significant variance in the number of CFU detected within media types relative to their means, often greater than the variation between media types. Importantly, PeptoneCG (no sugars) had fewer CFU than SabCG ‘stock’ (using pre-prepared complete powder) and ‘from scratch’ (using the same individual components used in other media), but comparable CFU to MaltosePeptoneCG and GlucoseMaltosePeptoneCG. The AgarCG and MaltoseCG were similarly very low in detected CFU, and each had fungal colonies that were similarly very poorly developed and difficult to see, in no way comparable to colonies observed on the other media, and very difficult to identify although several had *Alternaria*-like chains of dark spores at the surface despite a lack of distinct hyphae, hence definitely not the full gamut of outdoor airborne organisms, and would not normally be counted as CFU at 3 days incubation.

**Fig 1.**
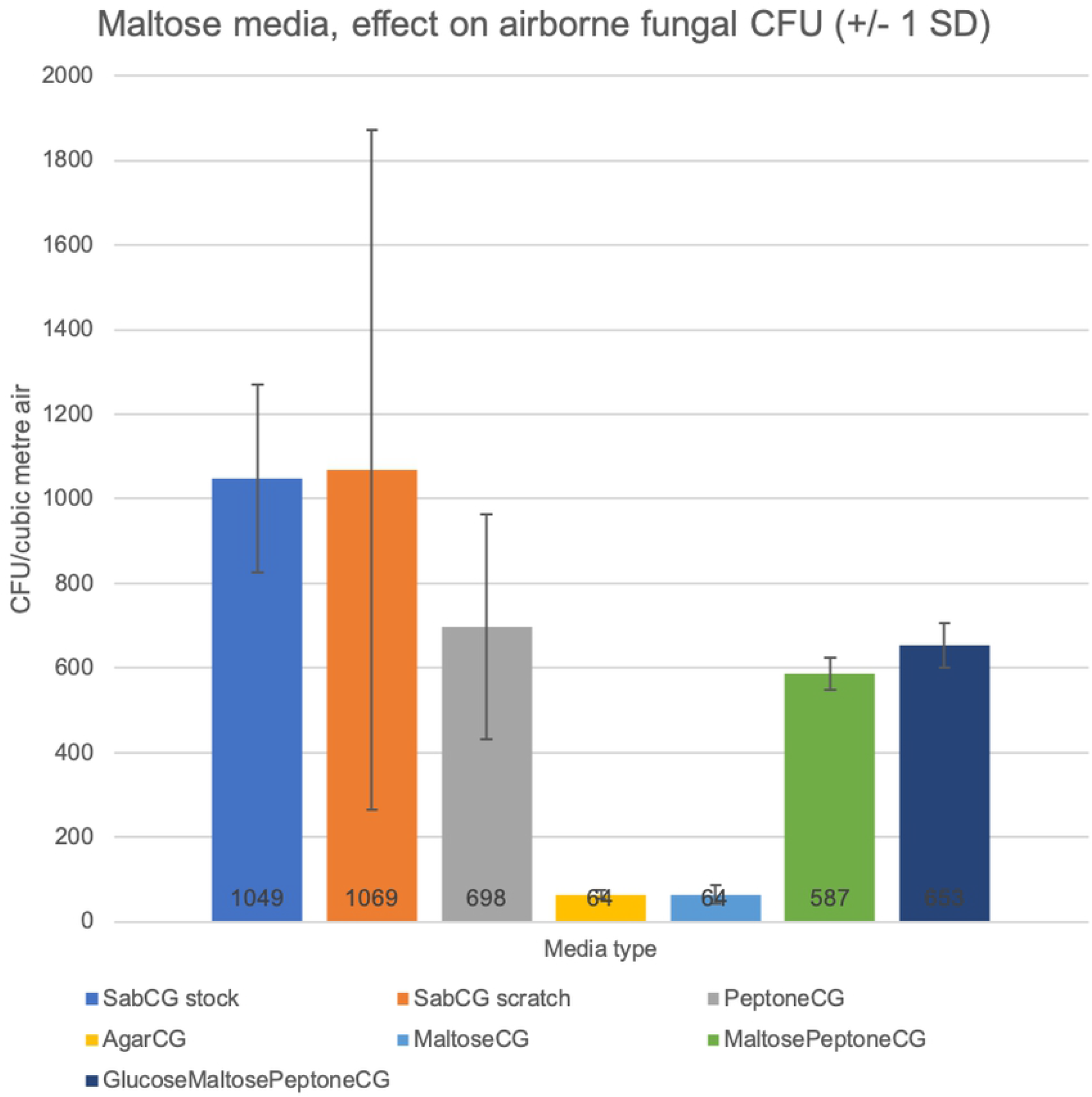
Maltose utilisation by outdoor airborne fungi. Media included SabCG ‘premade stock’ with CG antibiotics added, SabCG ‘scratch’ made from the individual components used in the other media, PeptoneCG (with no added sugars), AgarCG, 4% MaltosePeptoneCG and GlucoseMaltosePeptoneCG (hence 8% total sugars). Error bars are +/- 1 SD to better indicate the noted variance between three replicate plates of each medium.

One-way ANOVA analysis indicated was a significant difference between all media (p = 0.0089), but for SabCG(stock), SabCG(scratch) and PeptoneCG, significantly more variation within groups than the variation between groups (p = 0.623).

The results suggested that maltose was not commonly utilised by the fungi sampled from the outdoor air, being similar to PeptoneCG despite a noted degree of hydrolysis of maltose to glucose. The variation was quite significant and hence the association was not very strong at the number of replicate plates used per experiment.

That the PeptoneCG medium showed appreciable numbers of CFU was somewhat surprising as the standard understanding is that fungi typically require carbohydrates. The standard SabCG with 2-4% glucose seems OK for practical purposes, but the 4% (the standard formula of many years) is likely slightly more useful in comparisons with historical data.

That the GlucoseMaltosePeptoneCG was comparable to MaltosePeptoneCG and PeptoneCG was curious as it was expected to be similar to SabCG. It was noted that this would have been 8% sugars, double the standard 4% and hence possibly an effect of higher osmolarity, and/or catabolite repression.

MEA and maltose media without added glucose did have a detectable amount of glucose after autoclave sterilisation via test (Accu-Chek Mobile U1, Roche). Tests indicated 7.5 - 10.2 mM glucose in maltose-based media, and over 55-100 mM in MEA, and no detectable glucose in fresh maltose in cold RO water at approximately 100 mg/mL.

Hence there was a degree of hydrolysis of maltose into glucose likely during autoclave sterilisation at 121°C for 15 min, plus warm-up and cool-down time, and time at approx. 50-70°C during pouring. It was originally intended that the maltose solution be filter-sterilised and added to cooled liquid agar media, but this was not the case and instead this experiment was used mainly to demonstrate that when maltose is in a media, it does hydrolyse to glucose to a physiologically significant degree during normal autoclave sterilisation. The noted glucose concentration suggested approximately 1.8 g/L maltose had hydrolysed out of 40 g/L initial maltose, or 4.5%. It is also noted that normal human blood glucose concentration is approx. 5-10 mM. The presence of peptone was evidently the more critical factor for fungal growth, however.

### Clarified V8® Original juice, tomato juice, celery juice agar media supplements and heat-labile factor effects

There was no significant difference between media with or without various clarified vegetable juice extracts overlain over the top of SabCG media, and hence no significant heat labile factor nor missing vital vitamin or mineral or such supplied by these vegetables at least that enhances the detection of outdoor airborne fungi in mid-April (autumn), Melbourne, Australia (Figs 2 and 3). It was noted that uncooked tomato juice significantly changed odour when heated, going from a ‘grassy’ fragrance to a ‘tomato soup’ odour, yet this had no discernible effect on numbers of CFU nor colony morphology, range of cultured organisms, etc.

**Fig 2.**
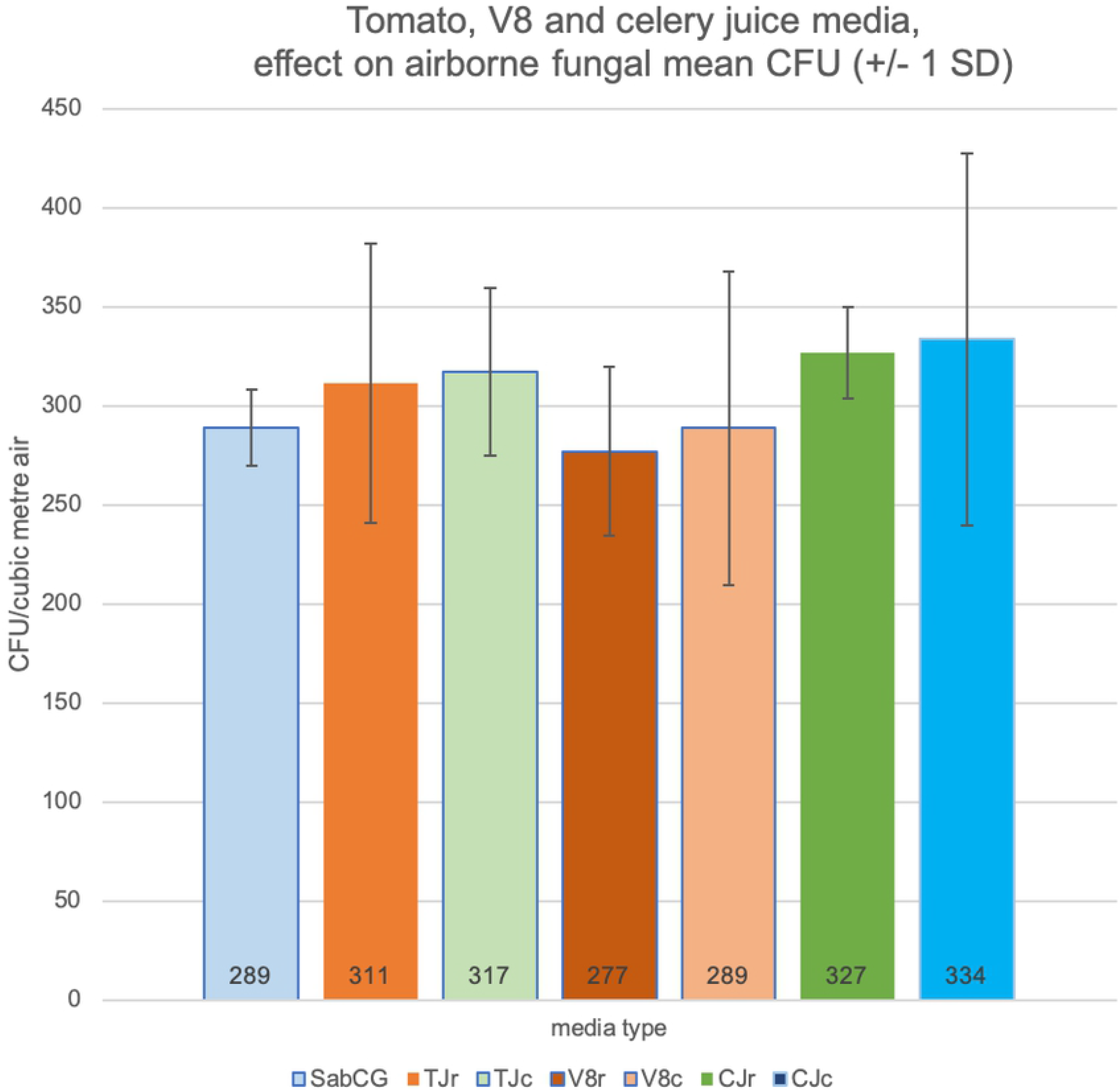
Media supplemented with raw or cooked vegetable juices. SabCG (1 mL water control); TJr / TJc = 1 mL Tomato Juice, raw / cooked; V8r / V8c = 1 mL V8^®^ Original juice, raw but supplied UHT pasteurised / cooked; CJr / CJc = 1 mL Celery Juice, raw / cooked. Each medium was tested in triplicate, and all were overlain over the top of pre-prepared SabCG media in Petri dishes.

**Fig 3.**
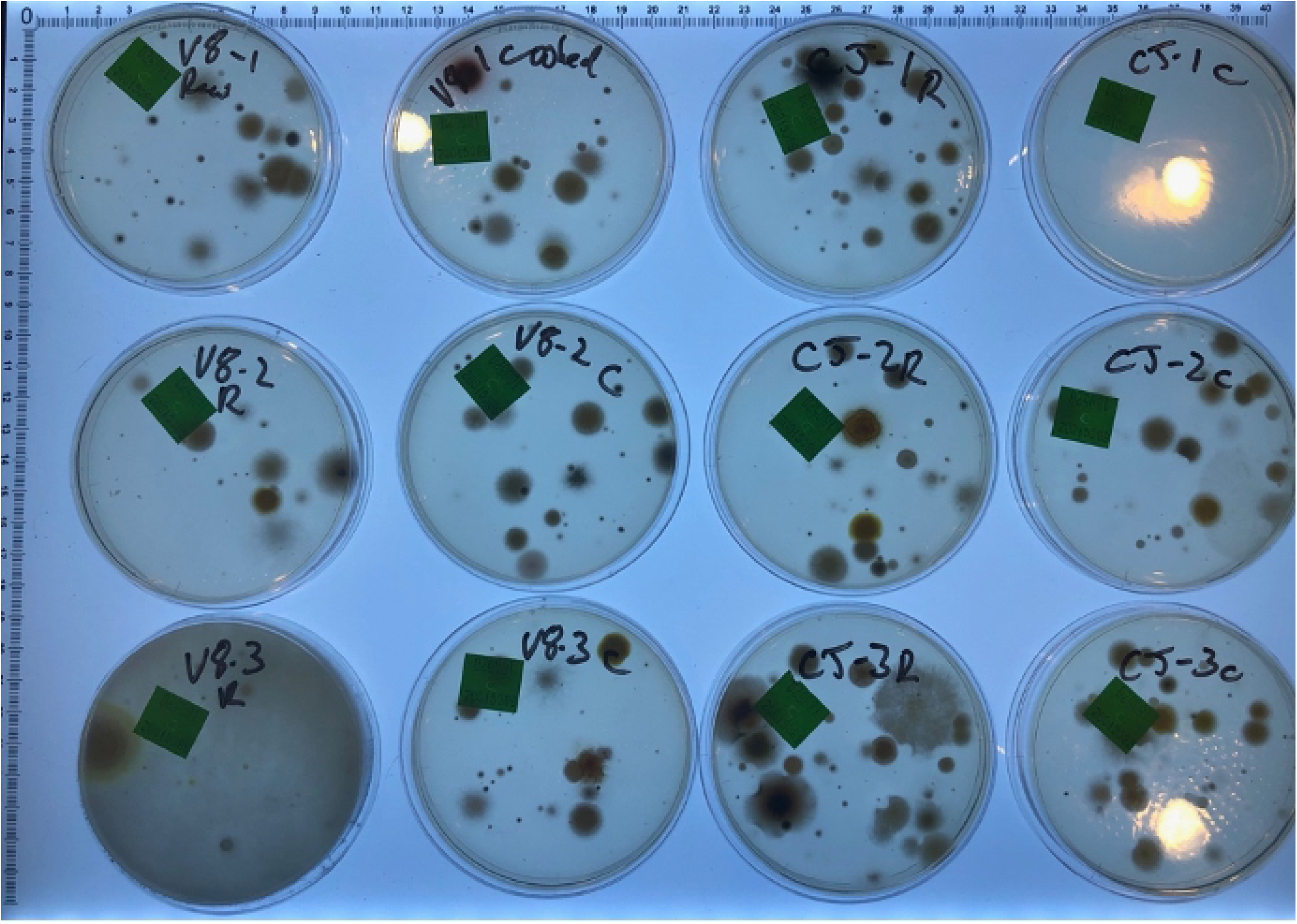
Images of results of raw and cooked vegetable juices. Images of some of the resulting Petri dishes (Fig 2) after culture for illustrative purposes only. Noted typical variation in numbers and types of fungi between each plate.

Early development cycles using vegetable juices that had not been clarified at all were found to not be useful for two reasons: being impossible to filter sterilise; being difficult to see through the tomato-based media from underneath, making enumeration and identification of colonies difficult. It was also determined by Lugol’s Iodine solution there was a significant amount of starch in unclarified juices that could presumably affect results by selectively advantaging organisms with amylase activity, further explored in other experiments presented below.

The method of adding 1 mL of clarified juices including V8^®^ Original juice over the top of SabCG media was useful but not equivalent to the media known as ‘V8 media,’ being approximately 1/3 strength and with glucose, peptone, chloramphenicol and gentamycin.

### V8c, DG18c and SabCG media

The V8c and DG18c media (containing antibiotics chloramphenicol and gentamycin, GC) were compared with SabCG via standard collection of airborne particles by 400-hole Andersen sampler outdoors in a suburban location on a winter’s afternoon with a light breeze (∼2 kn). Six replicate plates were used per medium, and collected in ‘collated’ sequence, being SabCG, DG18c, V8c, then repeating in that sequence to better allow for random changes in wind speed, direction and hence likely changes in numbers and types of viable airborne fungi over the course of the experiment, being approximately 2 h (Figs 4 and 5).

**Fig 4.**
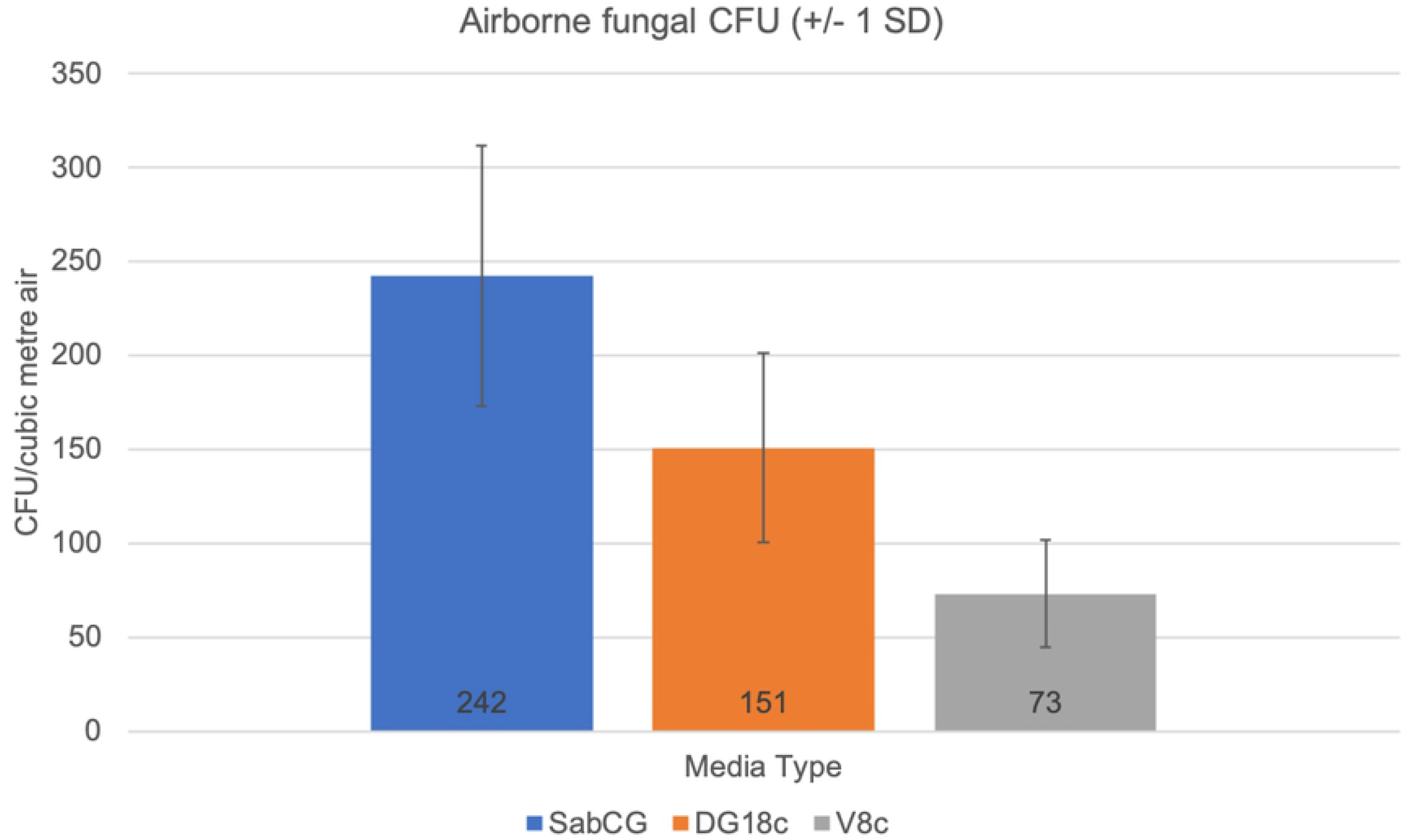
Graph of V8c, DG18c and SabCG media. 150 L outdoor air was sampled via Andersen 400-hole sampler onto six replicate plates of each of V8c, DG18c and SabCG agar media then incubated at 27°C for 3 days.

**Fig 5.**
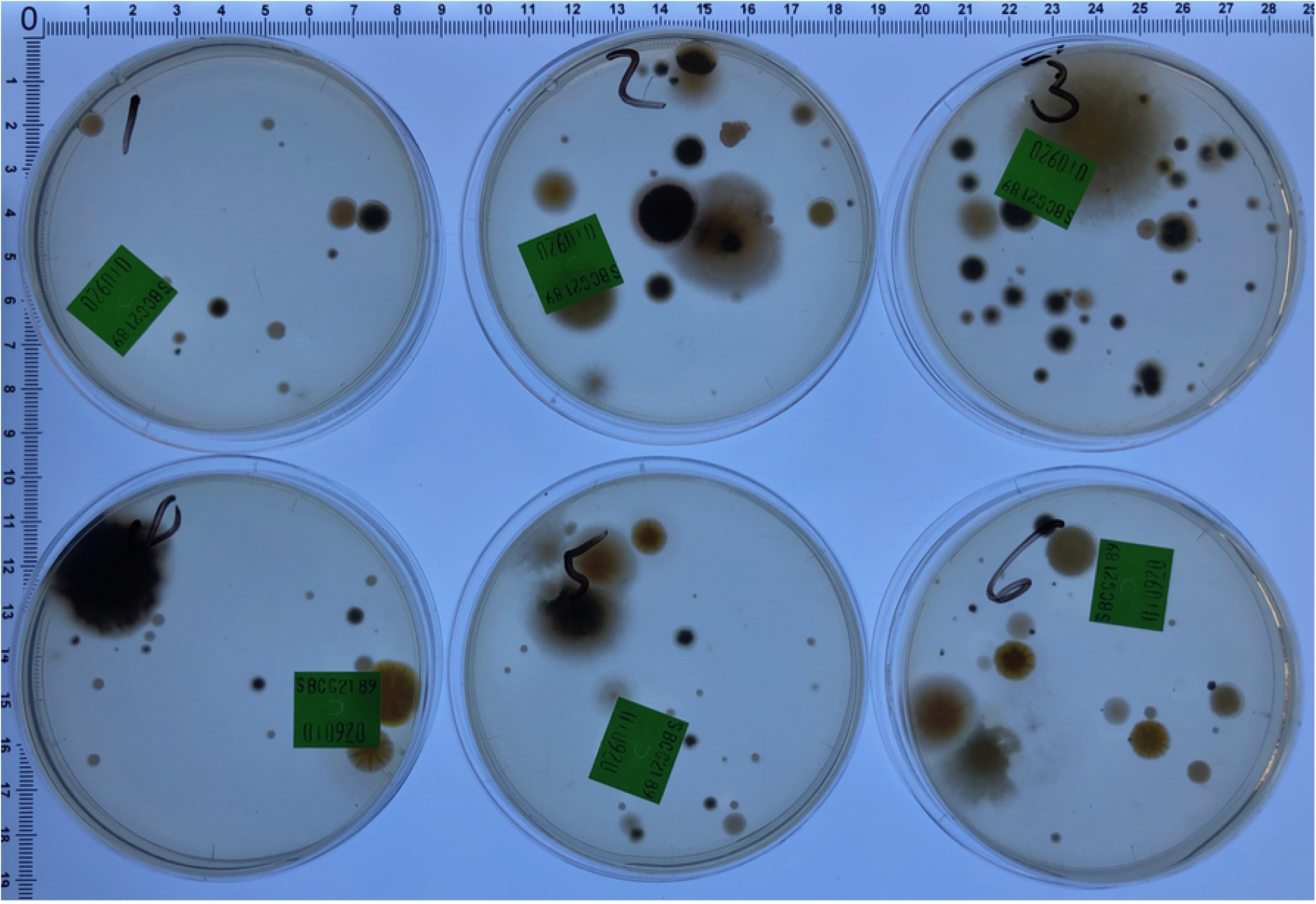
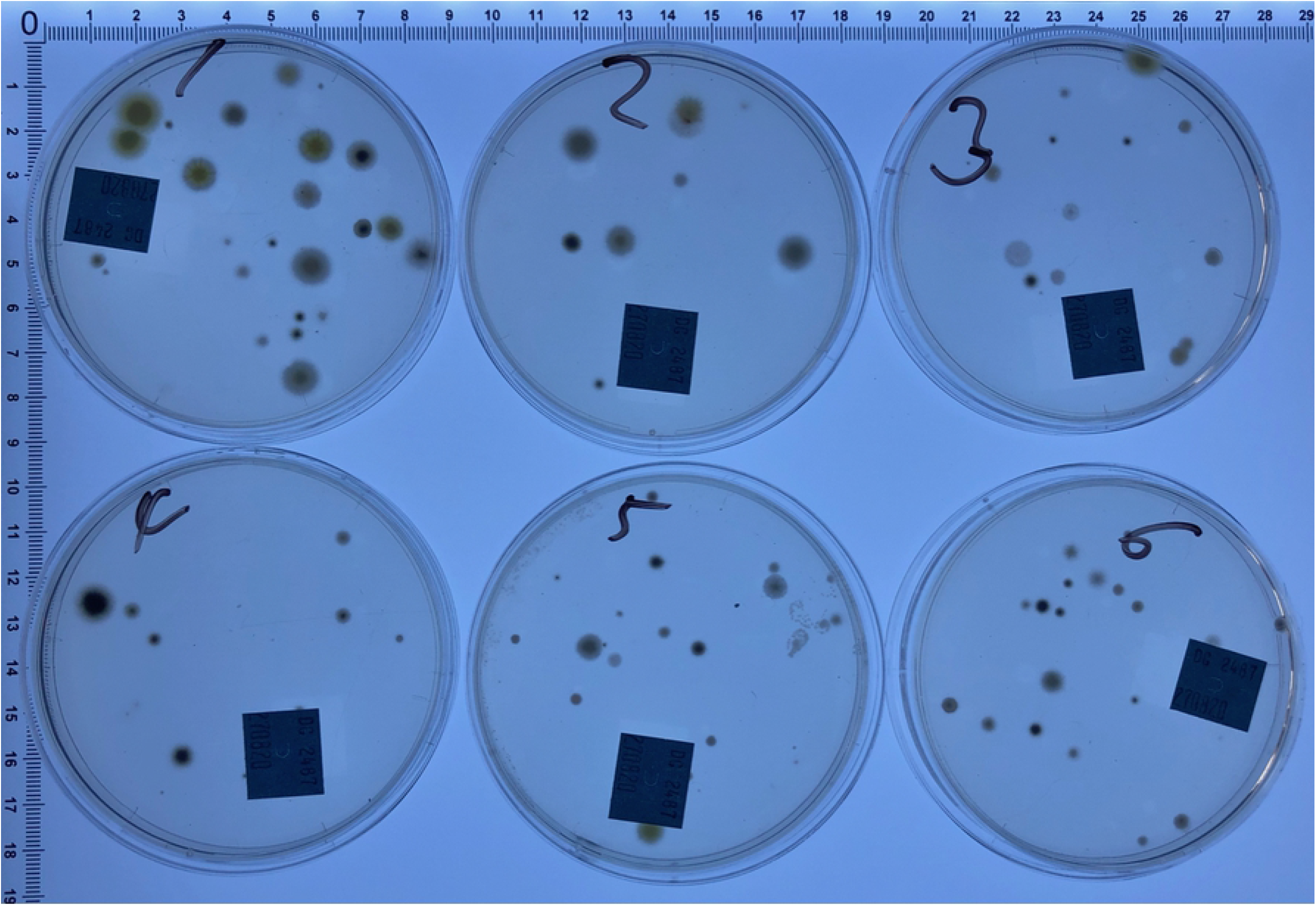
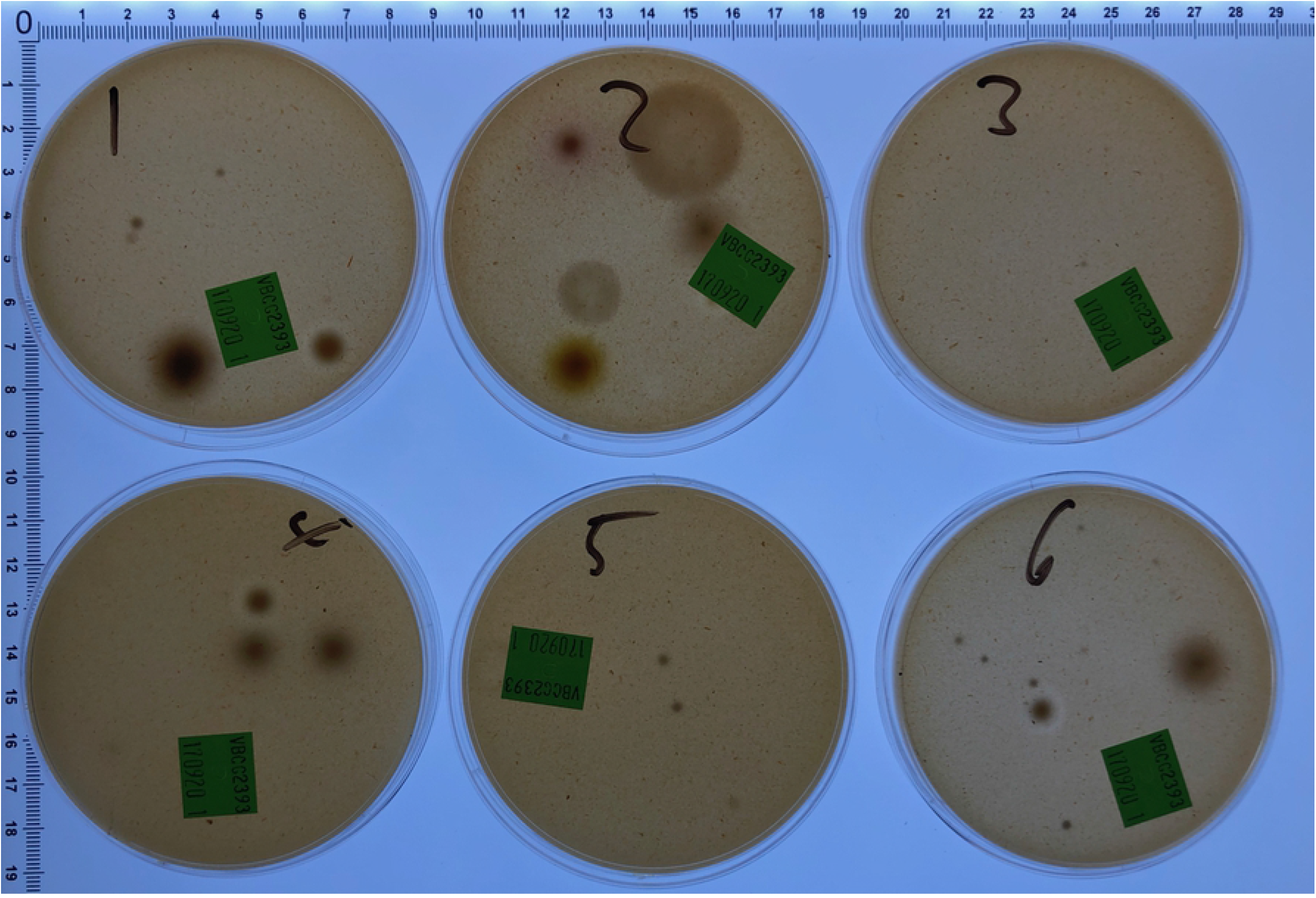
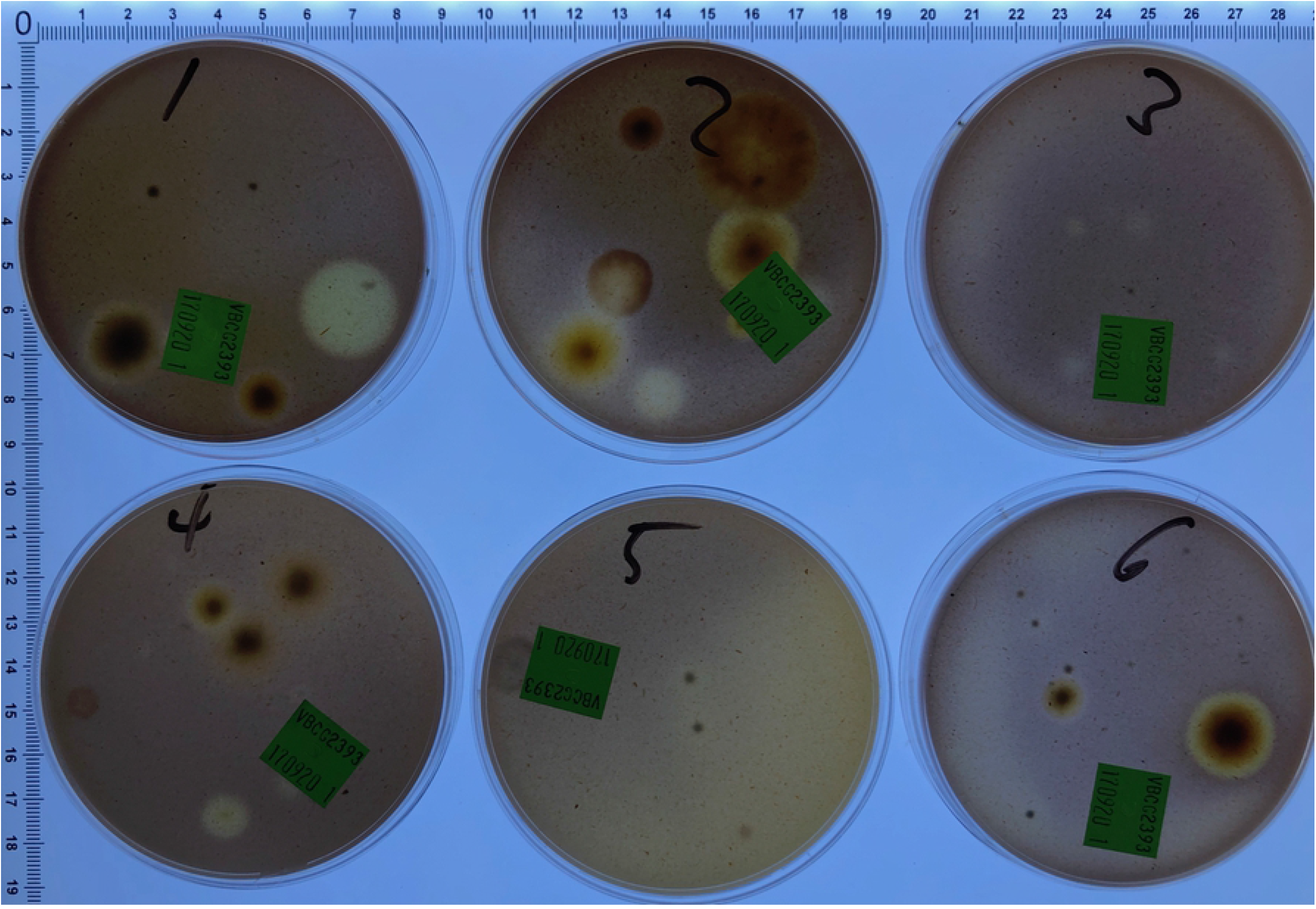
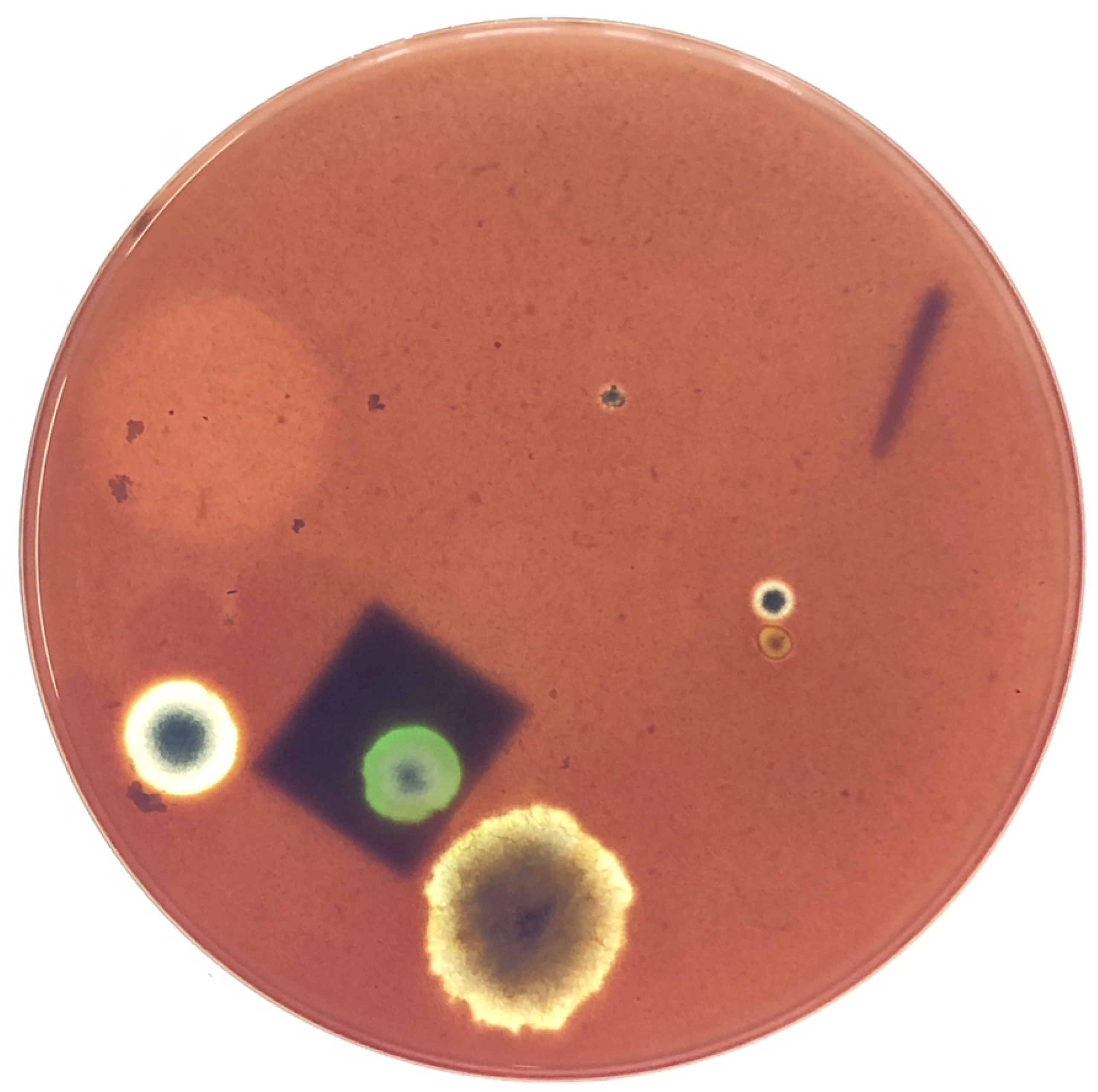
Images of SabCG, DG18c and V8c media. 150 L of outdoor air was sampled via Andersen 400-hole sampler onto six replicate plates of each of V8c (A), DG18c (B) and SabCG (C) agar media and incubated at 27°C for 3 days; (D) V8c media stained with Lugol’s Iodine, staining starch dark, seen from below the media (reverse side); (E) Detail of stained V8c media plate #1 from the upper side. Zones of clearing of starch was noted around some but not all colonies, with larger colonies being more associated with definite zones of clearing, smaller colonies without. Some seemingly colony-less zones of clearing may have been artefacts of wispy, low-mass colonies being rendered essentially invisible when the Lugol’s Iodine solution was added, causing structures to lay flat against the gel surface.

There was a significant difference between the three different media regarding the number of airborne CFU/m^3^ detected (P-value < 0.0003) when incubated for 3 days at 27°C.

The mean for SabCG was significantly highest at 242 CFU/m^3^, while DG18c and V8c were 151 and 73, respectively. Their corresponding SD were 69, 51 and 29, respectively, or 29%, 34% and 40% of their means, respectively. There was hence little overlap in their 68% confidence interval (i.e., +/- 1 SD around the mean), being 173 – 312, 100 – 201 and 45 – 102, respectively, assuming a normal distribution. The Kurtosis and skew of results for SabCG were 1.77 and 1.43, respectively, and for DG18c they were -1.89 and 0.66, respectively, and for V8c they were -0.54 and -0.03, respectively, and hence generally normal or only mild skew for DG18c and V8c, but significant for SabCG results, while Kurtosis was acceptable for each medium [71].

Additionally, the V8c medium was notably difficult to see through, being red in colour and nearly opaque and hence difficult to quickly observe the reverse / underside of many colonies, normally very useful in identifying/differentiating *Cladosporium* spp., c.f., *Aspergillus, Penicillium* spp.

It was also noted that the V8c medium had faint zones of clearing around some colonies but not all, and nearly always the colonies with clearing were large compared with colonies without zones of clearing (Fig 5E). This clearing was found to be due to the localised lack of starch in the media, as determined by flooding the plates with Lugol’s Iodine that stains starch dark, and hence likely due to digestion of the starch by some but not all organisms, and the organisms digesting starch growing more rapidly than those not doing so. The standard V8 agar medium formula does not include simple sugars such as glucose, nor peptone or similar alternative energy sources in any great abundance given V8® Original juice is stated as having 3.3 g/100 mL carbohydrates, of which 2.7 g/100 mL are sugars, 0.8 g/100 mL protein and 1.0 g/100 mL ‘dietary fibre.’

The DG18c medium did cause the colonies that grew to grow at a somewhat similar rate, and hence the colonies were more consistent in size at three days, but were quite often under-developed compared with SabCG, being without good maturation of spores, sporulating structures and typical colouration thus making identification more difficult / time-consuming.

It was also noted that the condensation from the DG18c was quite sticky, having significant amounts of glycerol presumably picked up while running over the gel that hence did not dry prior to, during or after sample collection. This caused one plate to become contaminated with a significant number of yeast colonies, causing it to be rejected from the data set. Typically, the condensate on the Petri dishes of other media without glycerol merely dry during the sampling of 150 L air, as also the media itself, typically visually apparent by the 400 dimples in the gel surface corresponding to the holes in the 400-hole Andersen sampler top-plate. The pattern of dimples is useful in determining that the bottom dish with media has not rotated during sampling due to vibrations from the air-pump as this affects the statistical calculations that are based on the assumption that a hole is either negative for growth (0 CFU), or has one or more viable CFU, and hence appears positive for growth despite possibly having multiple original viable CFU deposited on the gel surface.

It is unclear if the winter season may have caused a shift towards more high-a_w_-tolerant organisms, c.f., hot dry presumably lower a_w_ seasons, and hence higher apparent numbers in the high a_w_ SabCG medium cf. the lower a_w_ DG18c. Melbourne, Australia tends to have fairly dry, mild winters.

The use of six replicate plates was found to be useful (c.f., three) given the noted significant variation presumably due to the combination of the inherent uncertainty in the 400-hole Andersen collection method, and the uncertain nature of wind currents and weather. Repeating the experiment in other seasons is being considered as also during/after various weather events and conditions.

### Media tests: MEA, PDA with/without antibiotics, and mineral supplements MS1, MS2

There was no great difference in numbers of outdoor airborne fungi collected early-September (early spring) in Melbourne, Australia, between various mineral supplements and other media such as PDA or MEA, compared with SabCG (Figs 6 and 7). The variation between media groups was generally less than the variation within each group, but having the CG antibiotics present facilitated enumeration and identification of organisms.

**Fig 6.**
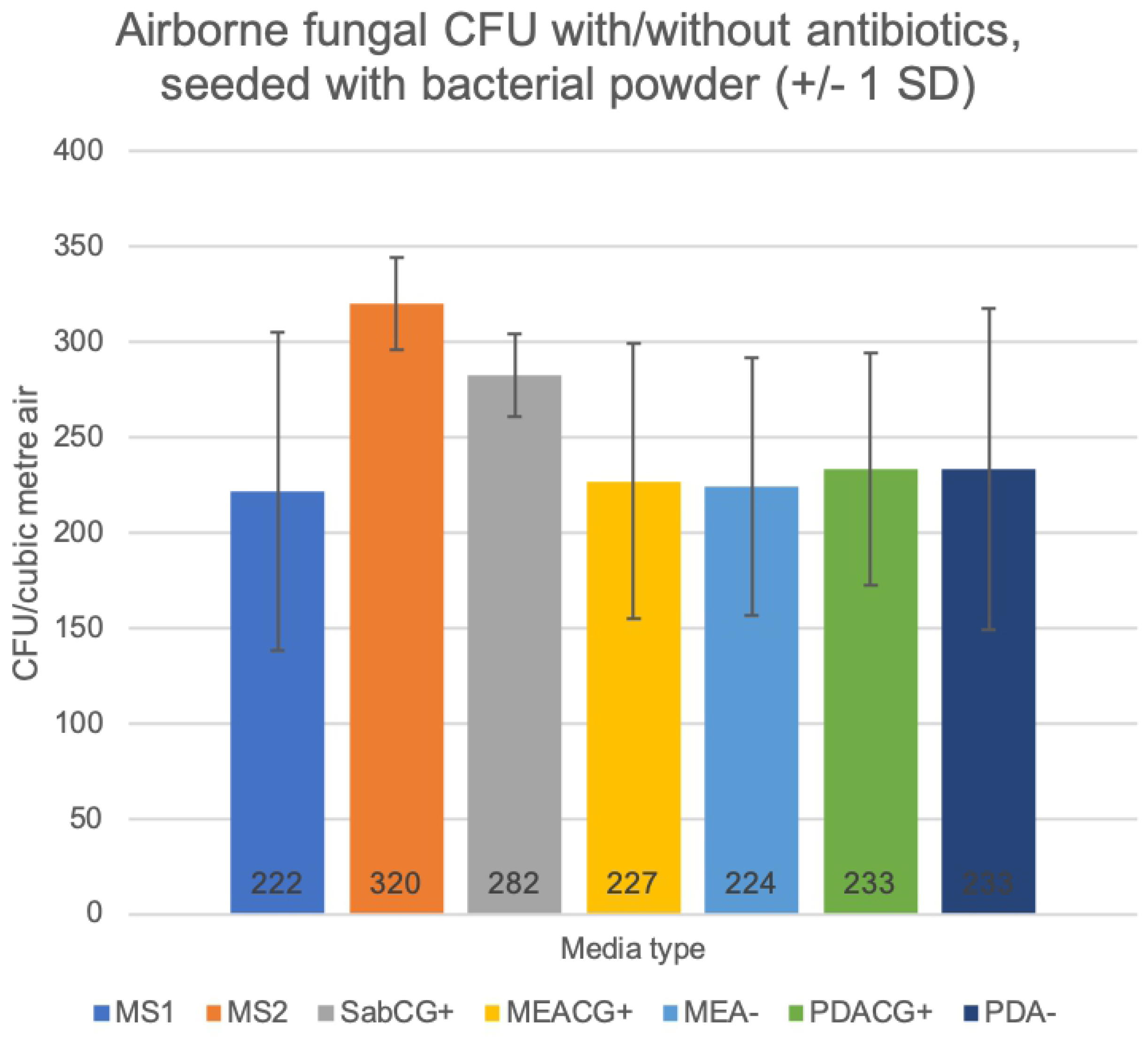
Effect of antibiotics in various media vs bacteria powder challenge. Mineral supplemented medias with CG antibiotics (MS1, MS2), other medias with antibiotics (SabCG+, MEACG+, PDACG+), and without antibiotics (MEA-, PDA-), seeded with airborne bacterial powder during sampling 150 L outdoor air, then incubated at 27°C for 3 days.

**Fig 7.**
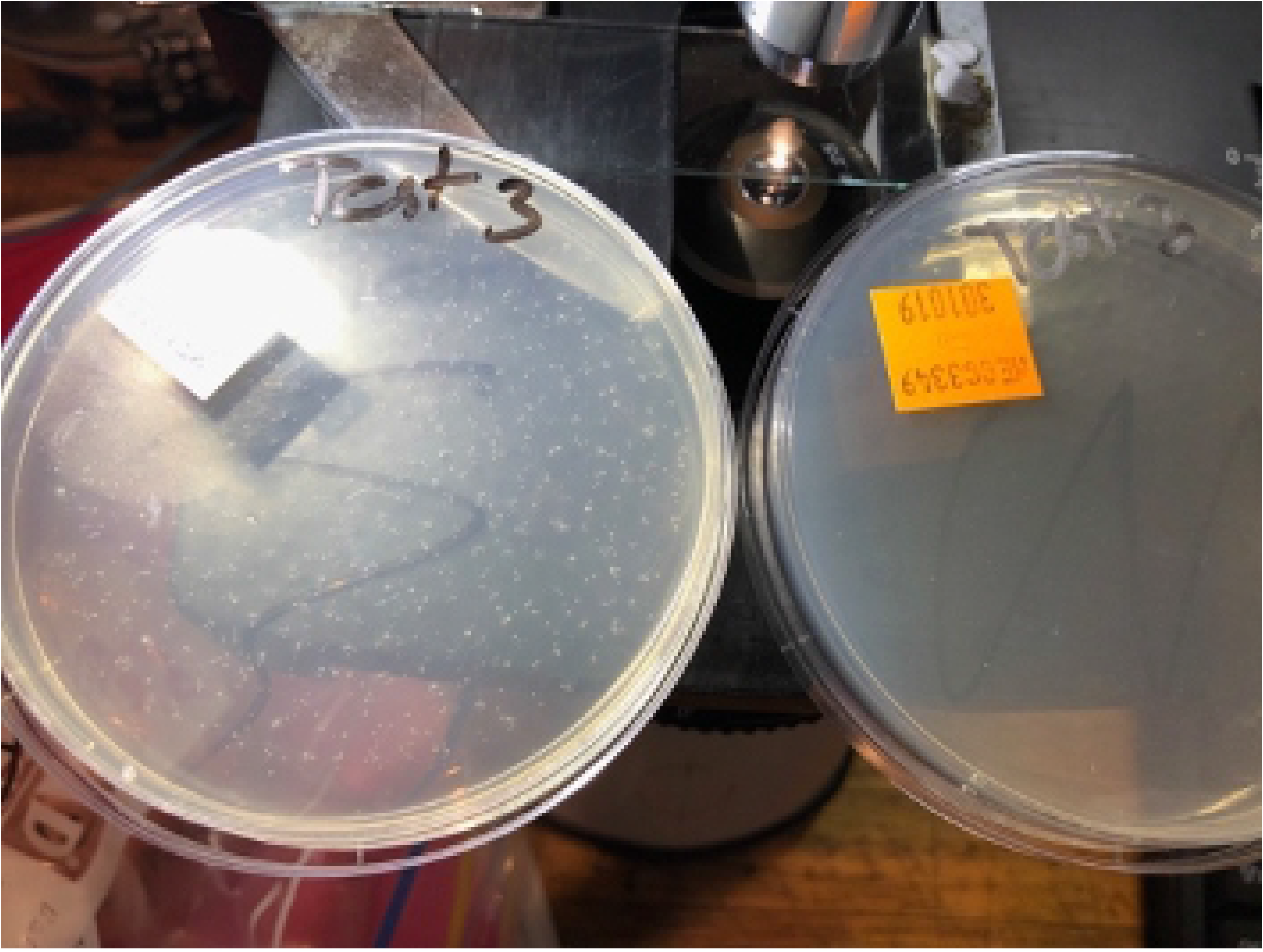
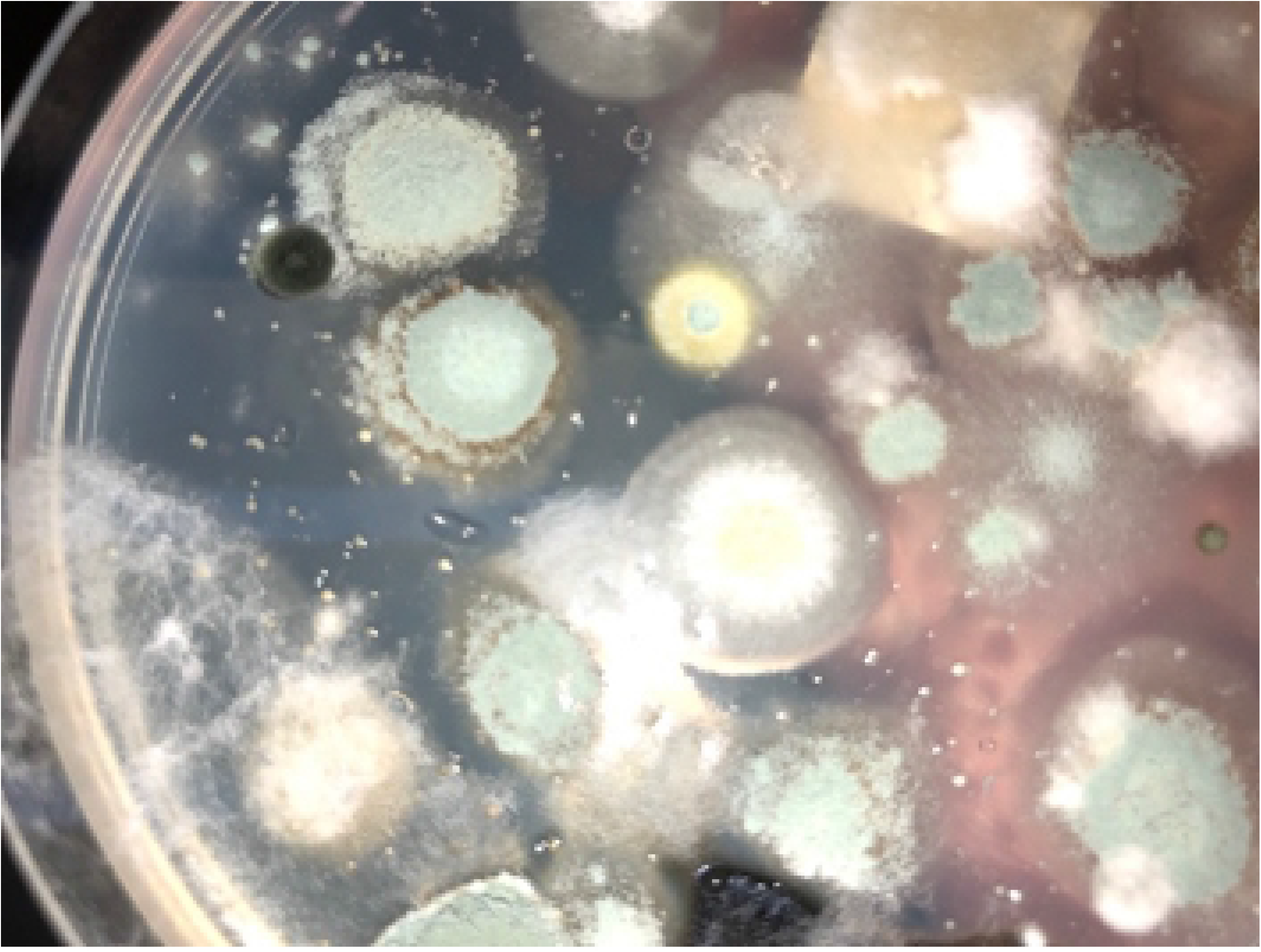
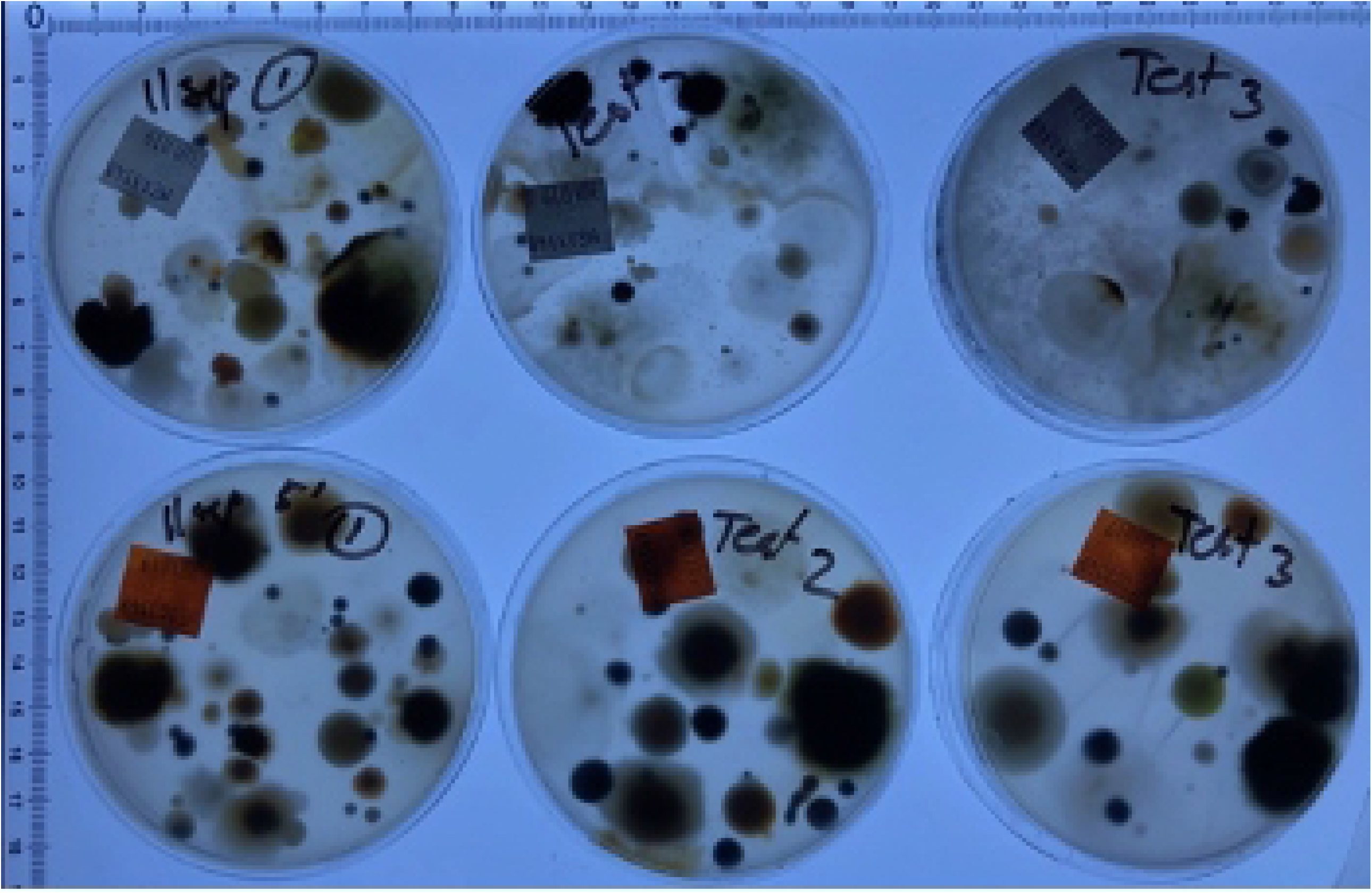
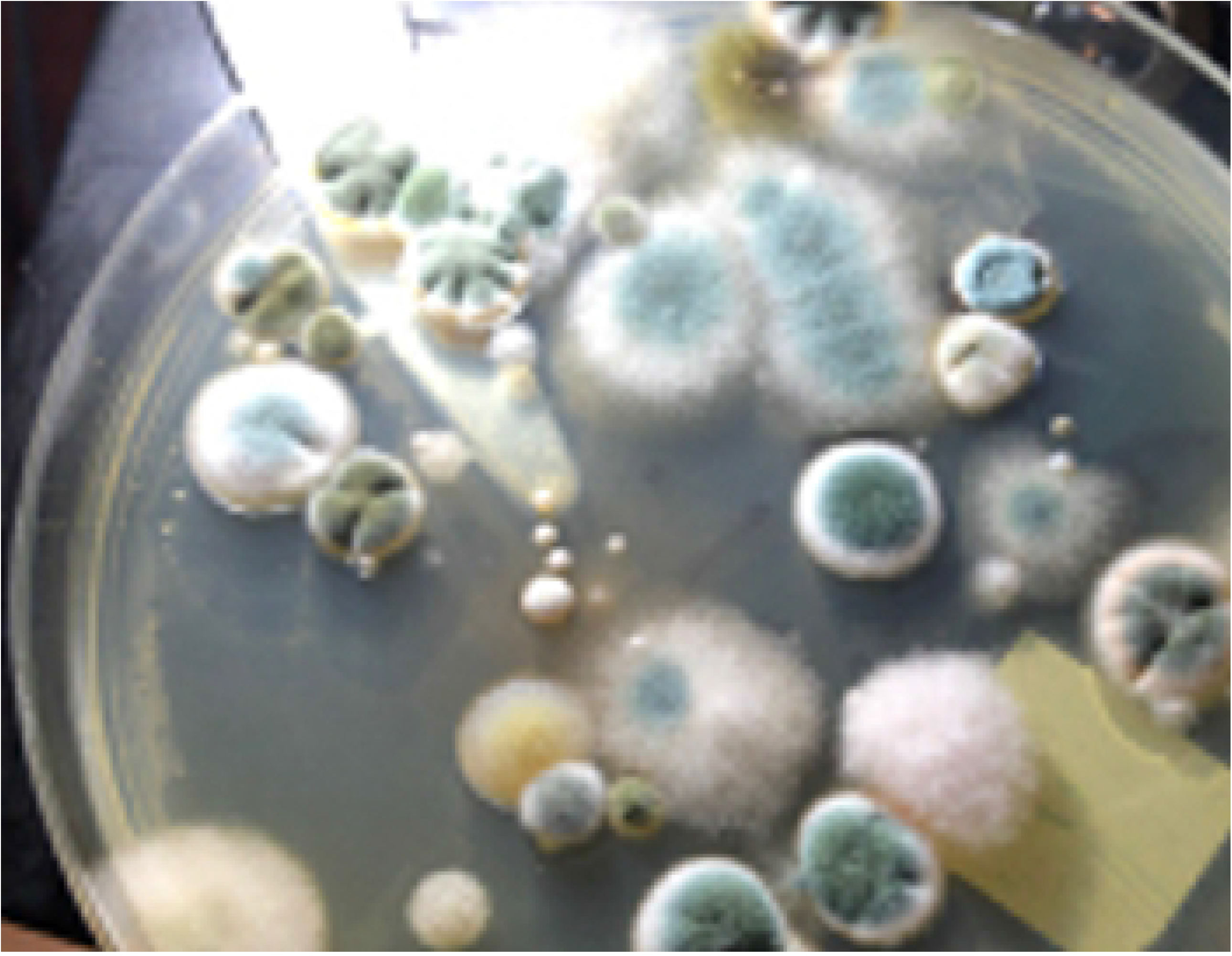
Visible effects of bacterial growth on fungi. Images of Petri dishes (Fig 6) seeded with bacteria powder while drawing 150 L outdoor air: (A) at 24 hr, 27°C, MEA (left) shows many bacterial colonies in the pattern of the 400-hole top-plate, and MEACG (right) showing no colonies due to CG antibiotics. Fungal colonies are not typically visible until 48 hrs, and not typically enumerated nor identified until 72 hr; (B) Example of MEA after 72 hr, 27°C, showing the small bacterial colonies in the 400-hole plate pattern. Fungal colonies were typically somewhat different to colonies on MEACG, having visibly different morphologies and colours, e.g., *Penicillium* spp. colonies were typically flatter, smaller, with scalloped edges and the spores were pale and at a delayed state of growth/development, while other fungi remained very sparse, spreading without apparently sporulating and hence making identification very difficult; (C) Example of results at 72 hr, 27°C, with MEA (top row) and MEACG (bottom row), with notably different colony colours and morphologies with/without CG antibiotics; (D) Example of typical colony morphologies on MEACG and other media with antibiotics such as SabCG, having better maturation, conidia development, colouration and overall more consistent colony shape thus aiding identification and enumeration.

Chloramphenicol / Gentamycin (CG) was useful in reducing numbers of bacterial colonies to zero, increasing the confidence in the identification of yeast colonies that usually look similar (shiny, glabrous, usually small, round colonies; Fig 7). Also the fungal colonies were more regular in shape, tending to have a rounded circumference, cf. irregular colonies with ‘holes’ and scalloping from bacterial colonies growing where the fungal colony would otherwise be, and fungal colonies having more regular colours, appearance, sporulation/fruiting bodies, etc., presumably due to not having to actively respond to competing bacteria nearby, or passively via the drain on available resources in the local media.

Mineral supplements MS1 and MS2 caused the media to go cloudy, which was less than ideal for counting and identification purposes. This cloudiness occasionally lessened over time and/or occasionally when colonies grew nearby, forming halos of clear areas presumably due to changes in pH due to atmospheric CO_2_ and/or biological processes and fermentation products also including CO_2_, and also likely ammonia, organic acids, etc.

### Various glucose concentrations of media vs airborne fungal detection

There is some small degree of difference in numbers of CFU (and their colony morphology) between SabCG media with different concentrations of glucose, with possibly a more ideal concentration being about 2% cf. the standard 4% (Figs 8 and 9). The effect is small, however, and would be unlikely to significantly influence counting or identification. There is little effect over the tested range from 1% to 8%, as predicted from the estimated a_w_. There was some effect noted in the peptone-CG-only medium, but even then it seems the majority of airborne fungi able to grow at 27°C within 3 days are able to substantially grow and sporulate without a sugar source, using the peptone as an energy and nitrogen source to complete their life-cycle, at least when sampled in early-February (summer), Melbourne, Australia. Glucose seemed to generally be of benefit, but some experiments suggested a possible suppression of growth at high sugar concentrations, initially hypothesised to be a catabolite-repression and/or high osmolarity / low a_w_ effect.

**Fig 8.**
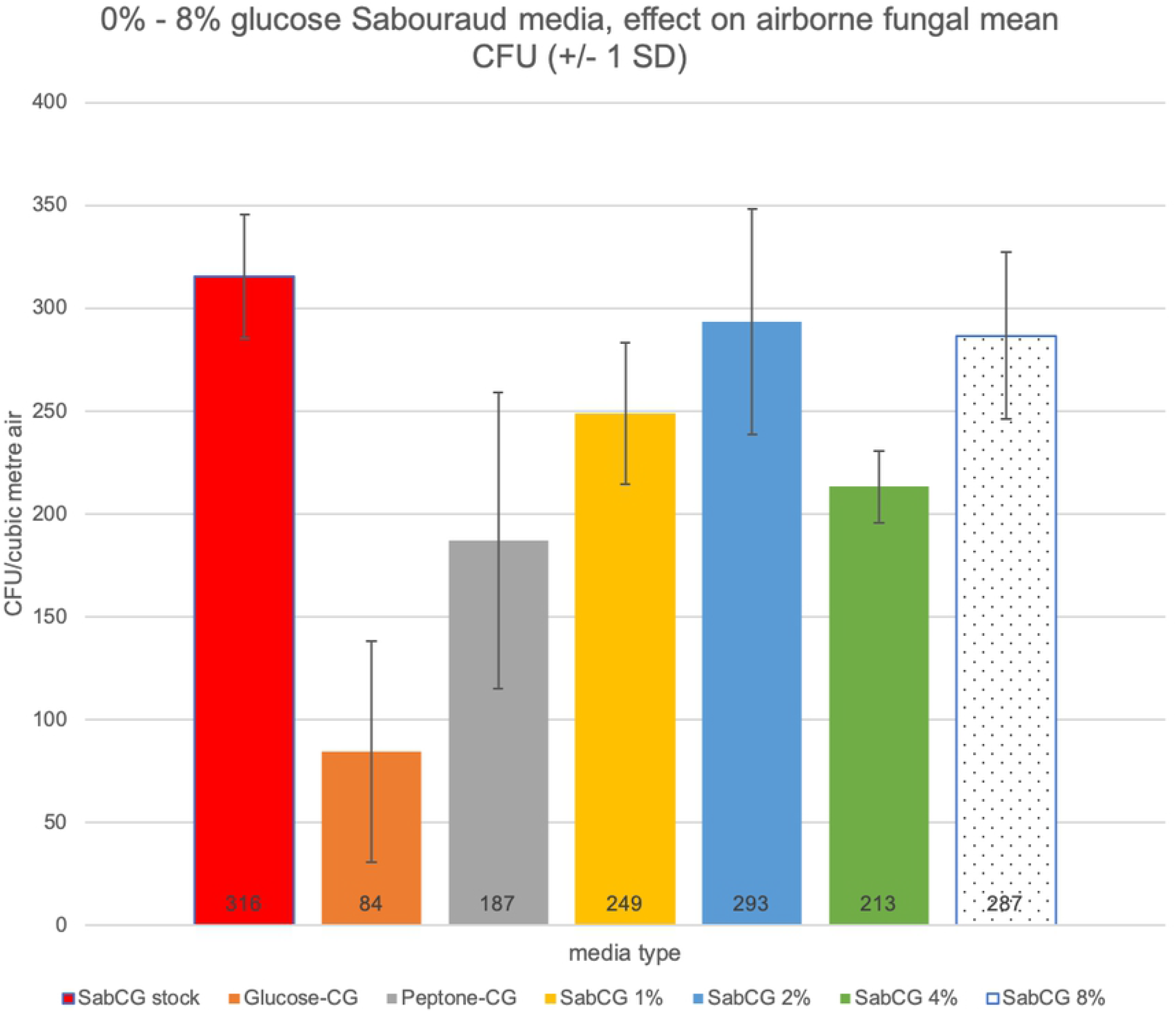
Various media glucose concentrations graph. Graph of CFU/m^3^ air (from 150 L air sampled) vs media with various glucose concentrations. SabCG stock is 4% glucose. The colonies on the Glucose-CG (no peptone, 4% glucose) medium were very under-developed and not strictly comparable with those on other media. Colonies on Peptone-CG were similar to those on SabCG 1%, 2%, 4% and 8% glucose.

**Fig 9.**
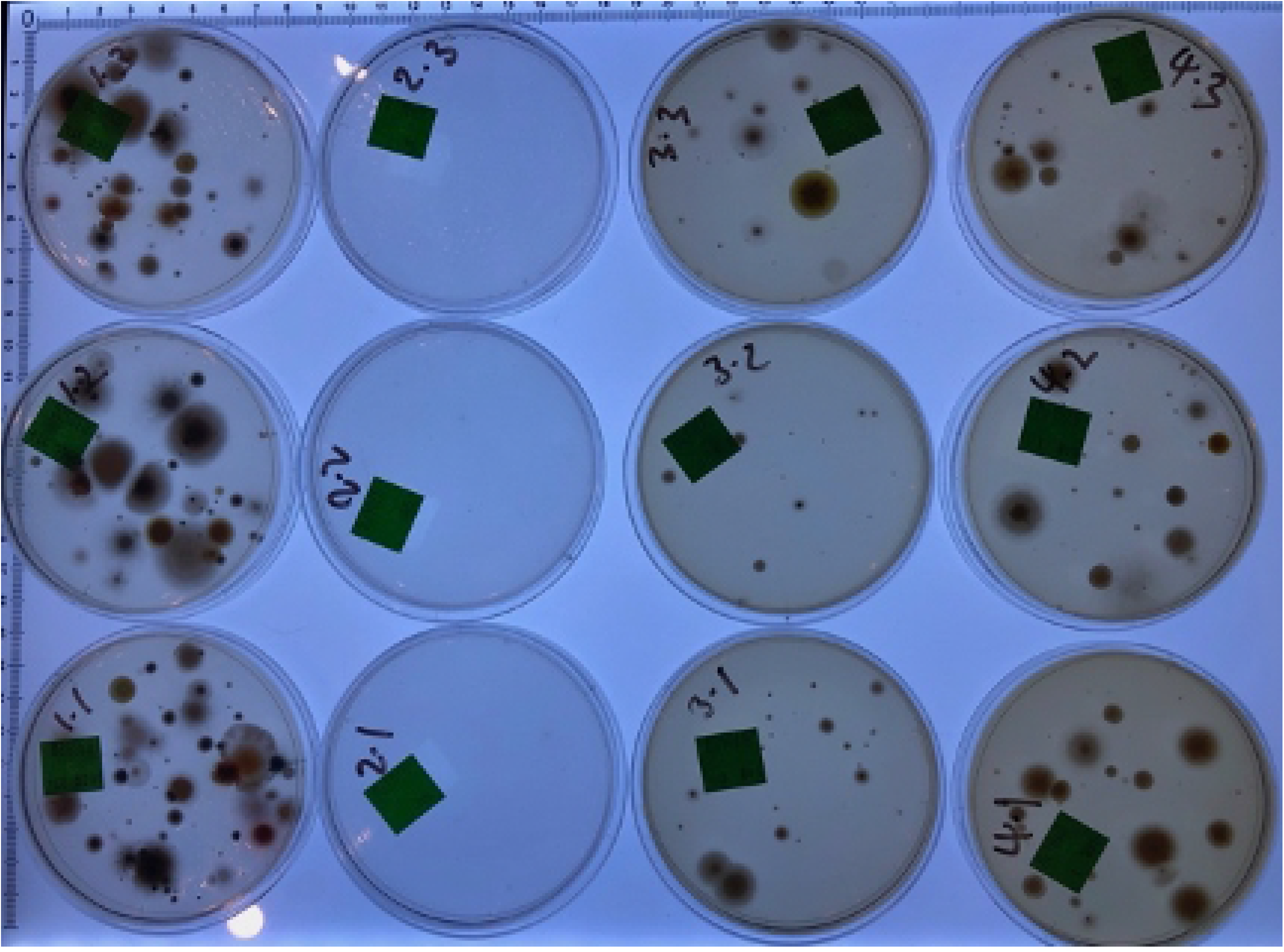
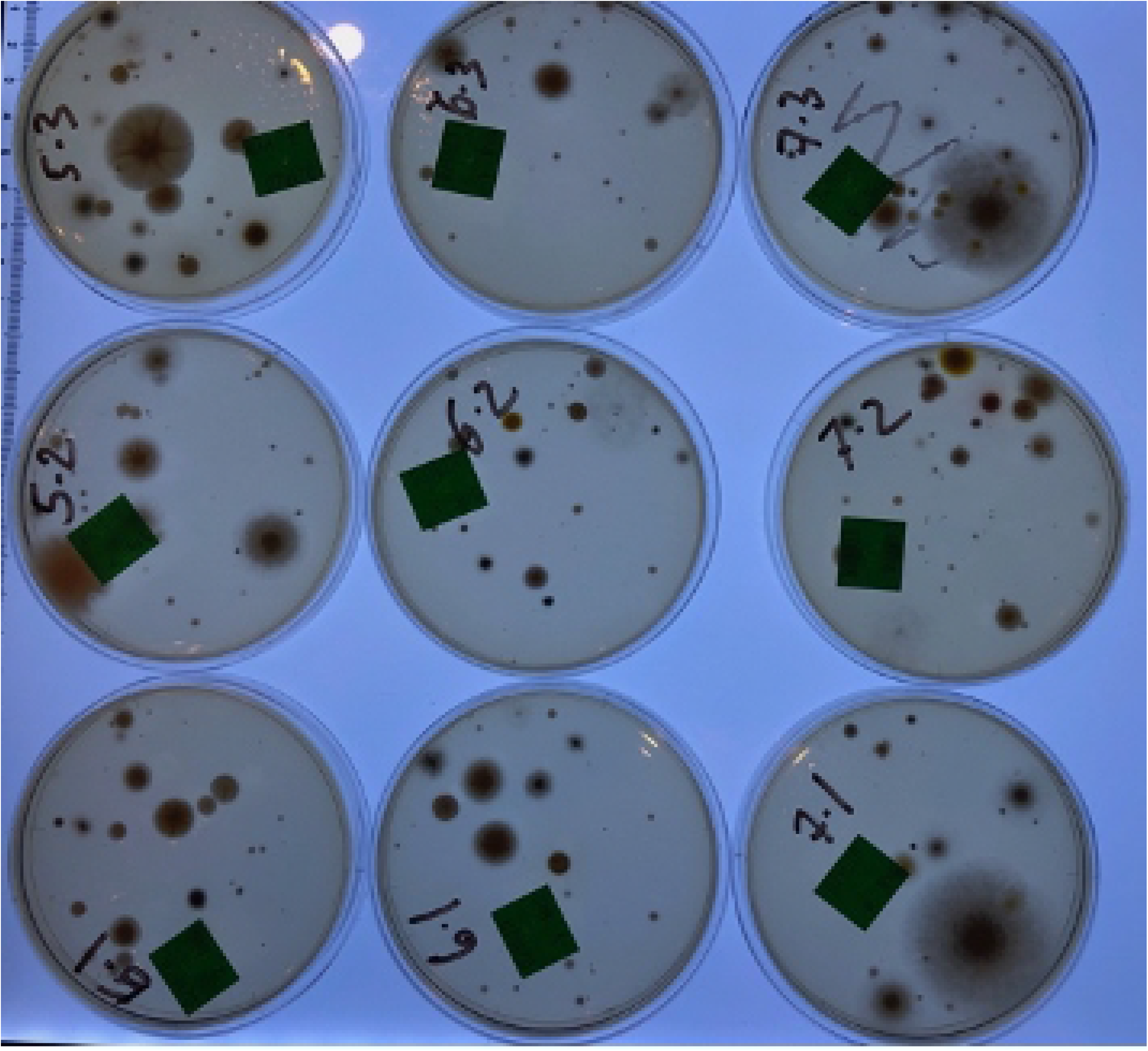
Images results of various glucose concentrations. Images of Petri dishes after culture as used in Fig 8: (A) Column at leftmost, Stock SabCG; second left, glucose-CG (colonies virtually invisible and very under-developed), third-left, peptone-CG, fourth-left / rightmost, SabCG 1% glucose; (B) Column at leftmost, SabCG 2% glucose, second-left, SabCG 4% glucose, third-left / rightmost, SabCG 8% glucose.

### Flame sterilisation/sanitisation of 400-hole Andersen air sampling top-plate

Dousing the aluminium Andersen 400-hole air sampling top-plate with alcohol and flaming it did effectively sterilise or at least sanitise it of significant numbers of dry viable *Penicillium* spores placed there (Fig 10). This was not entirely expected because it was presumed the heat would be insufficient to raise the temperature of the metal above that required for significant killing for long enough to do so. It is known that aluminium has a high thermal conductivity and is often used in heat-sinks and cookware. It was also noted that some charred debris and possibly inorganic grit was often left behind, however, which seemed to accumulate in the quite narrow holes, reducing air flow and the number of ‘open holes,’ critical to the operation of the Andersen 400-hole sampler.

**Fig 10.**
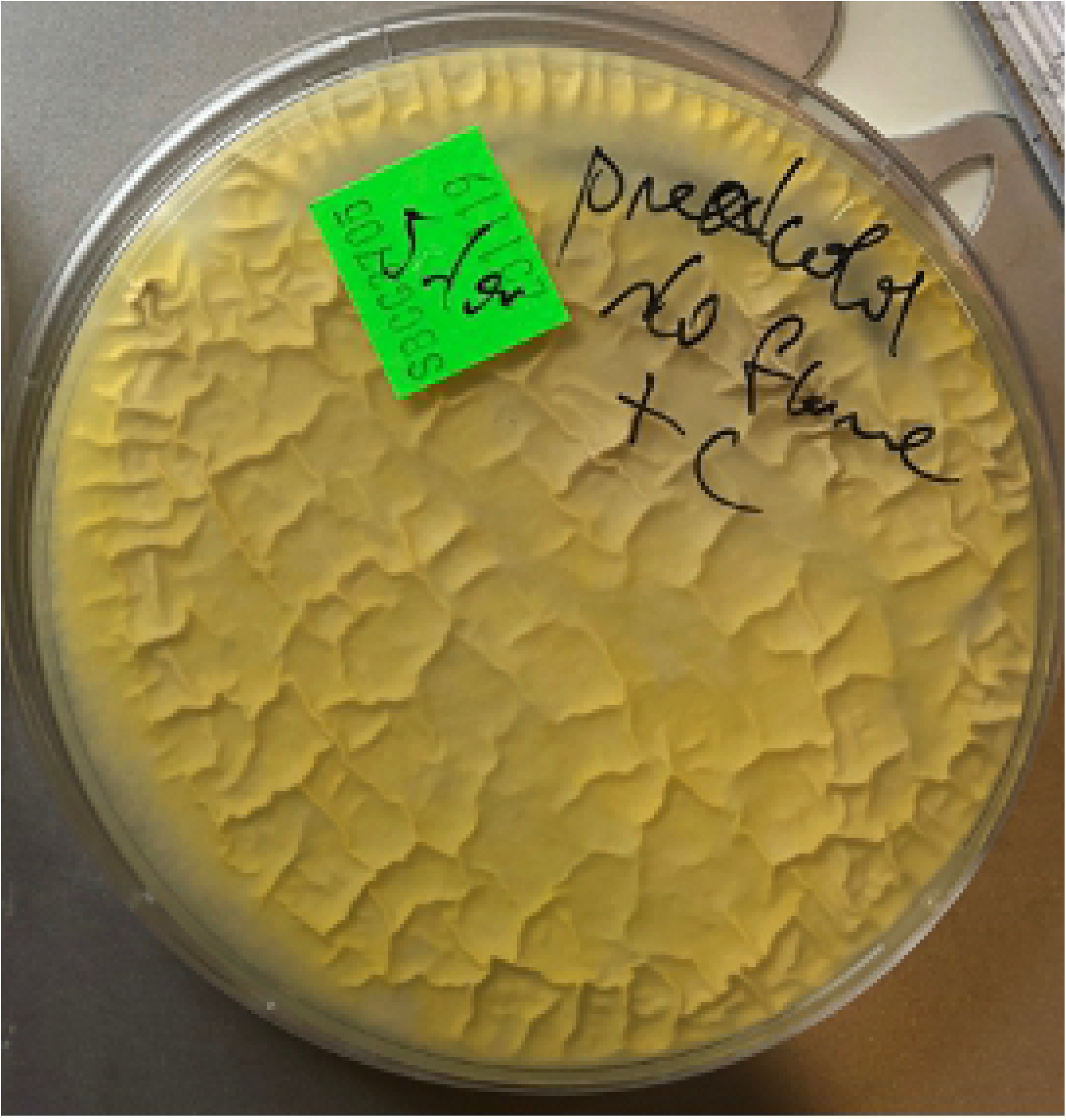
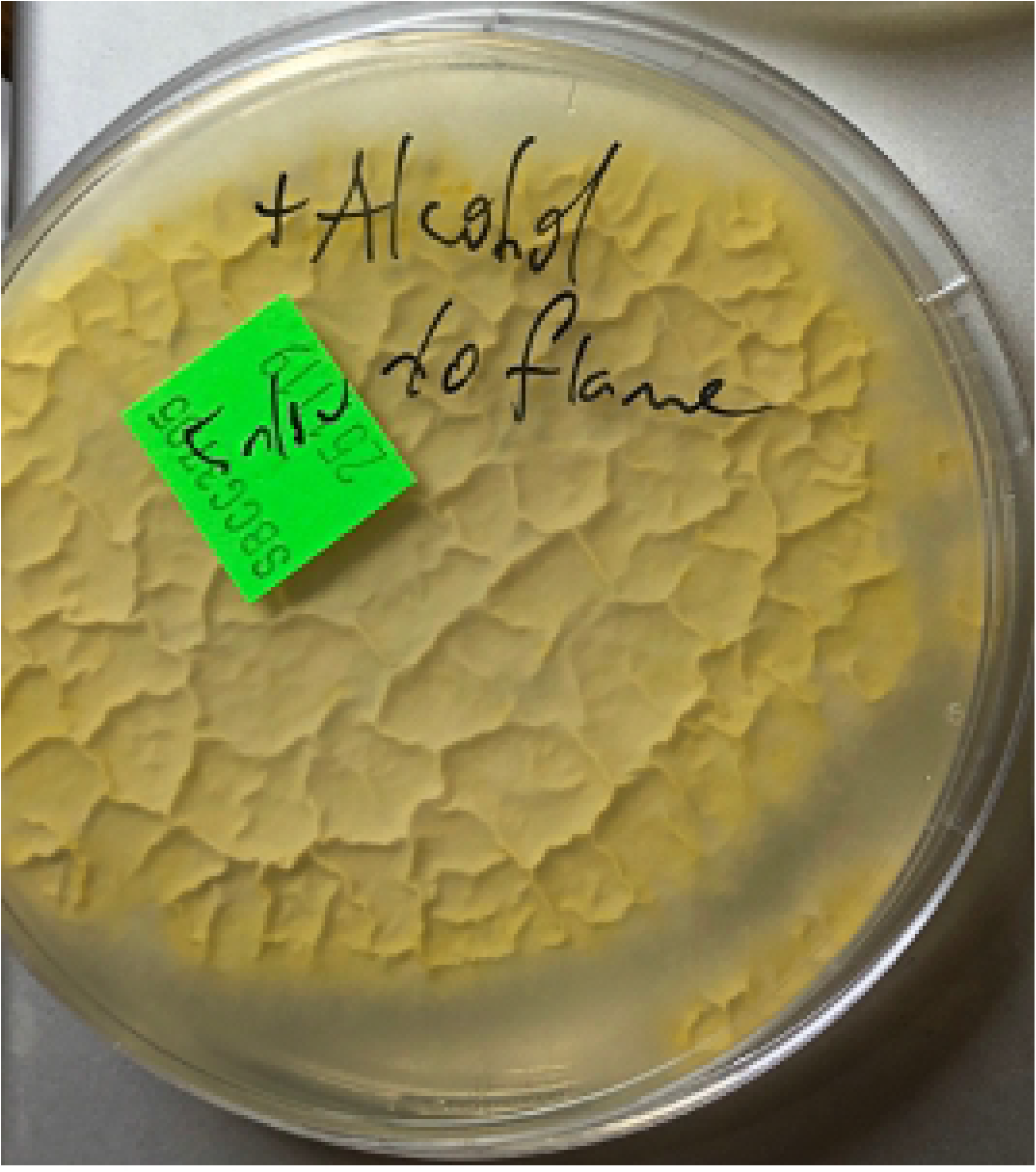
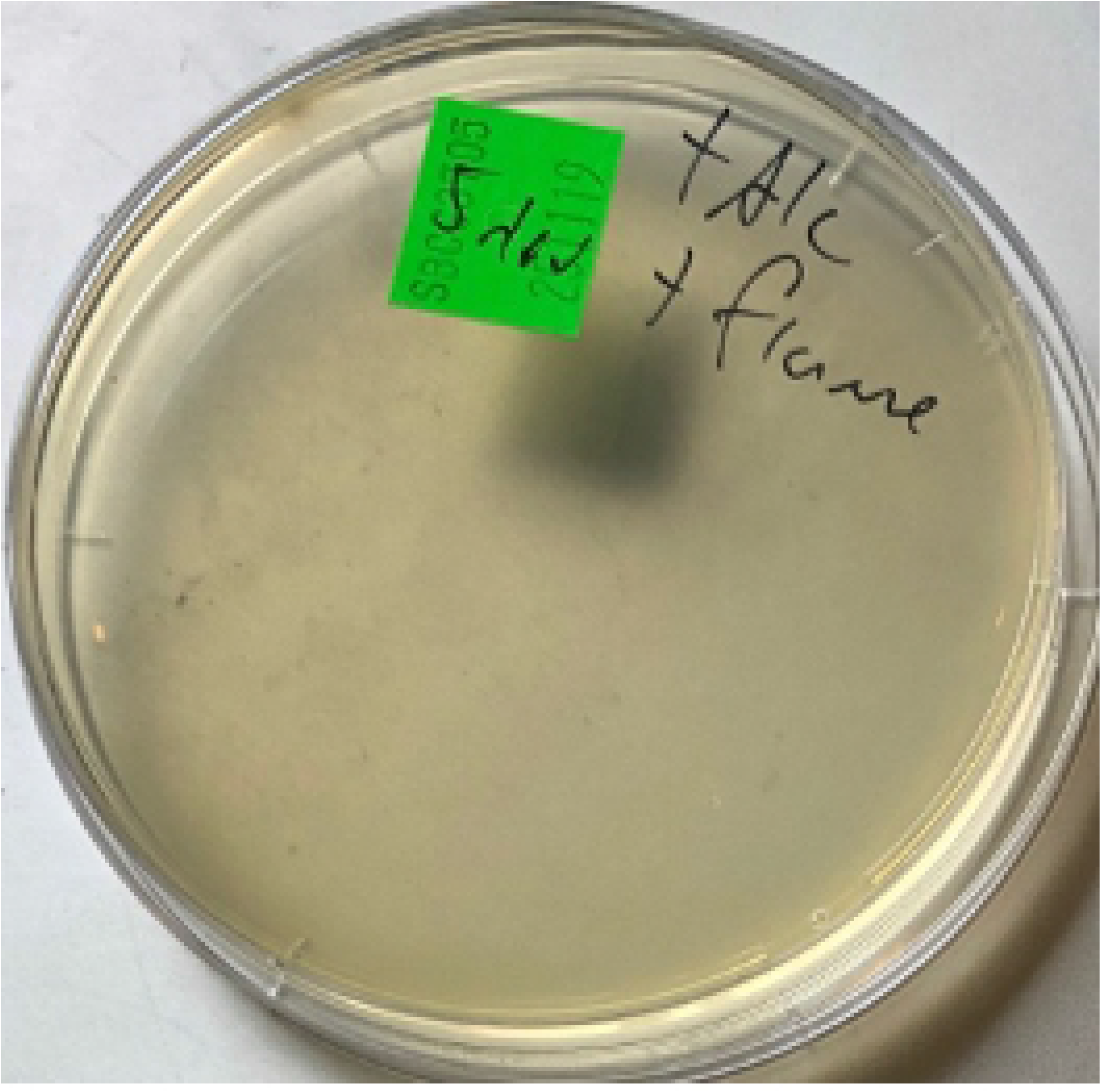
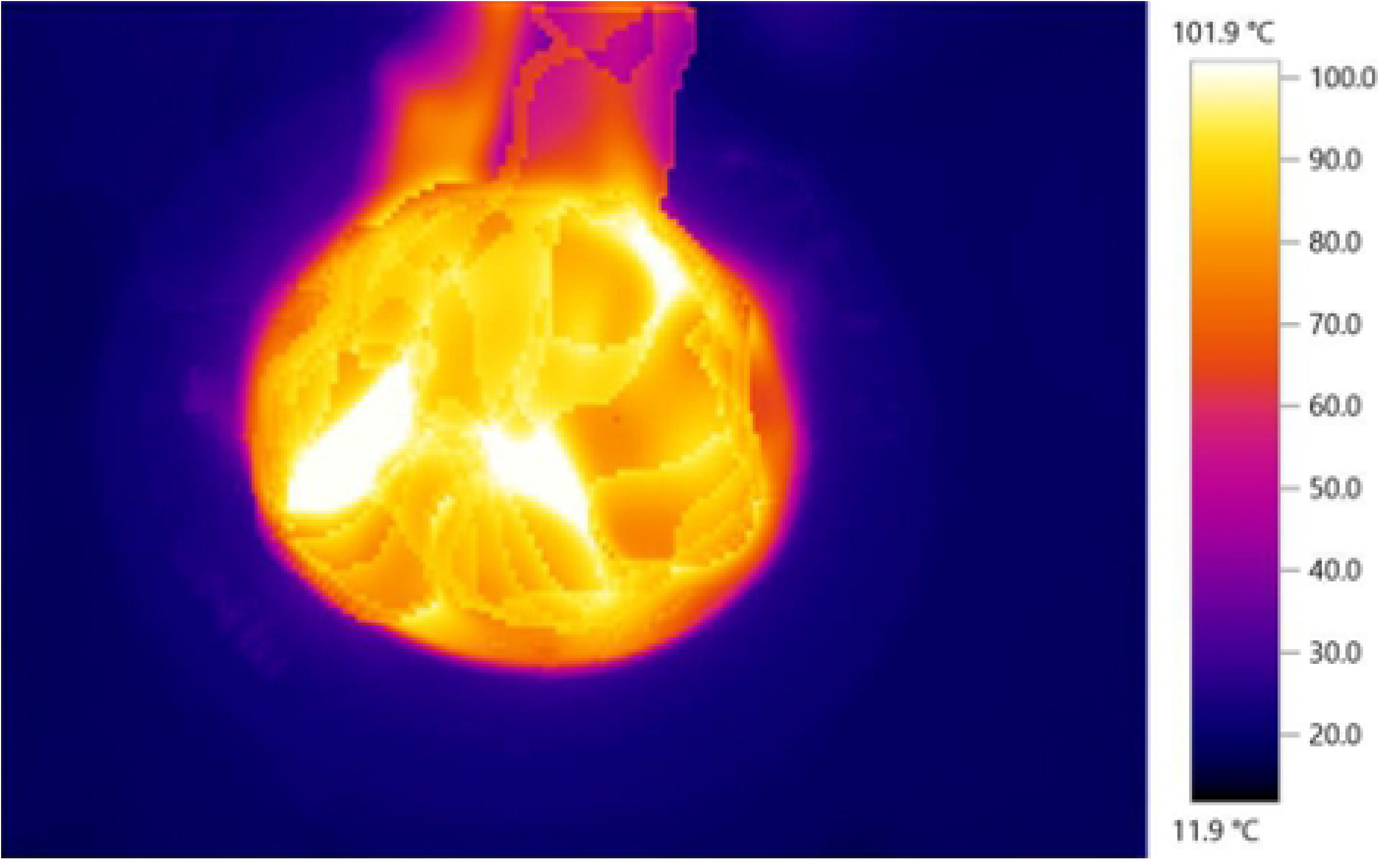
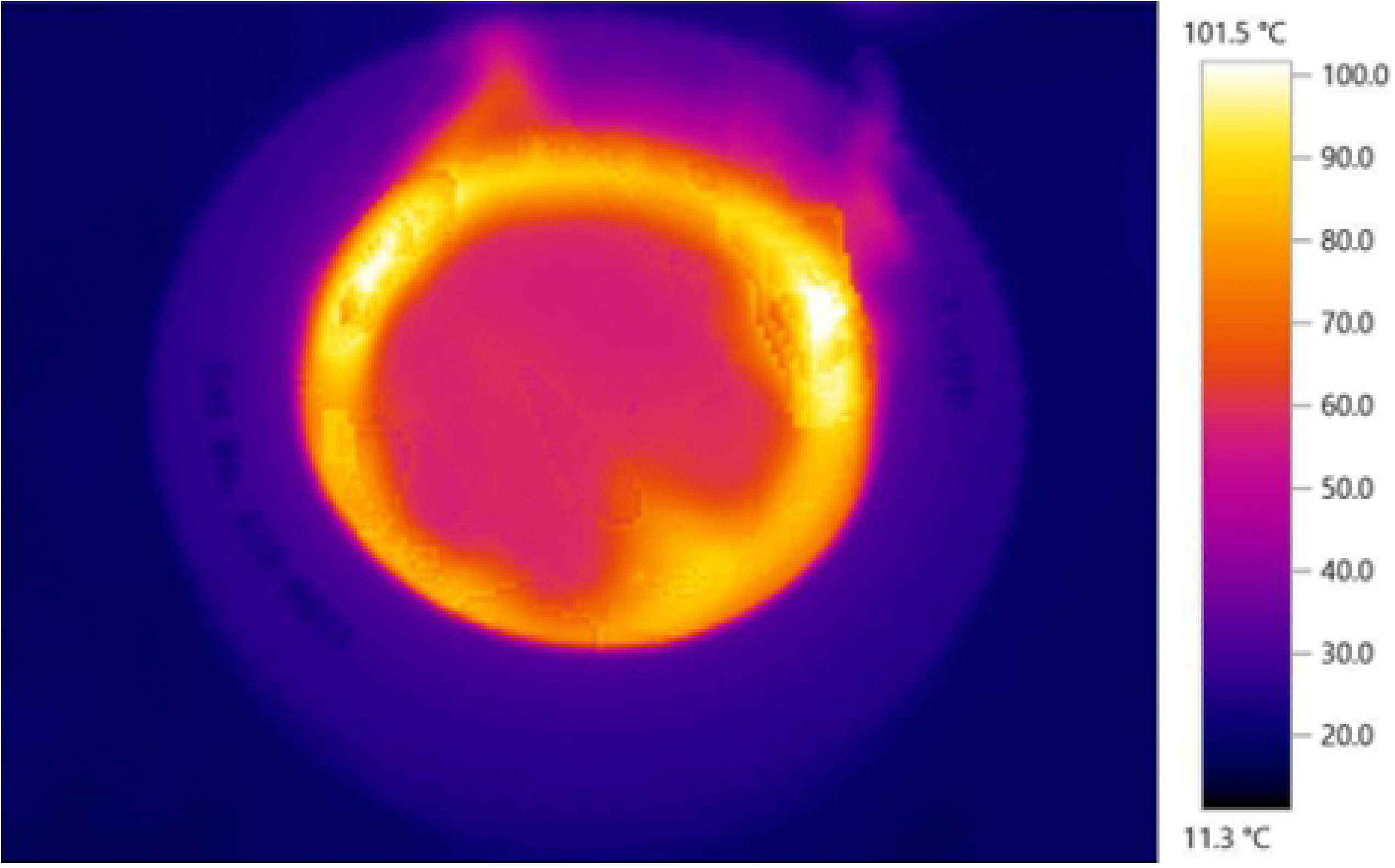
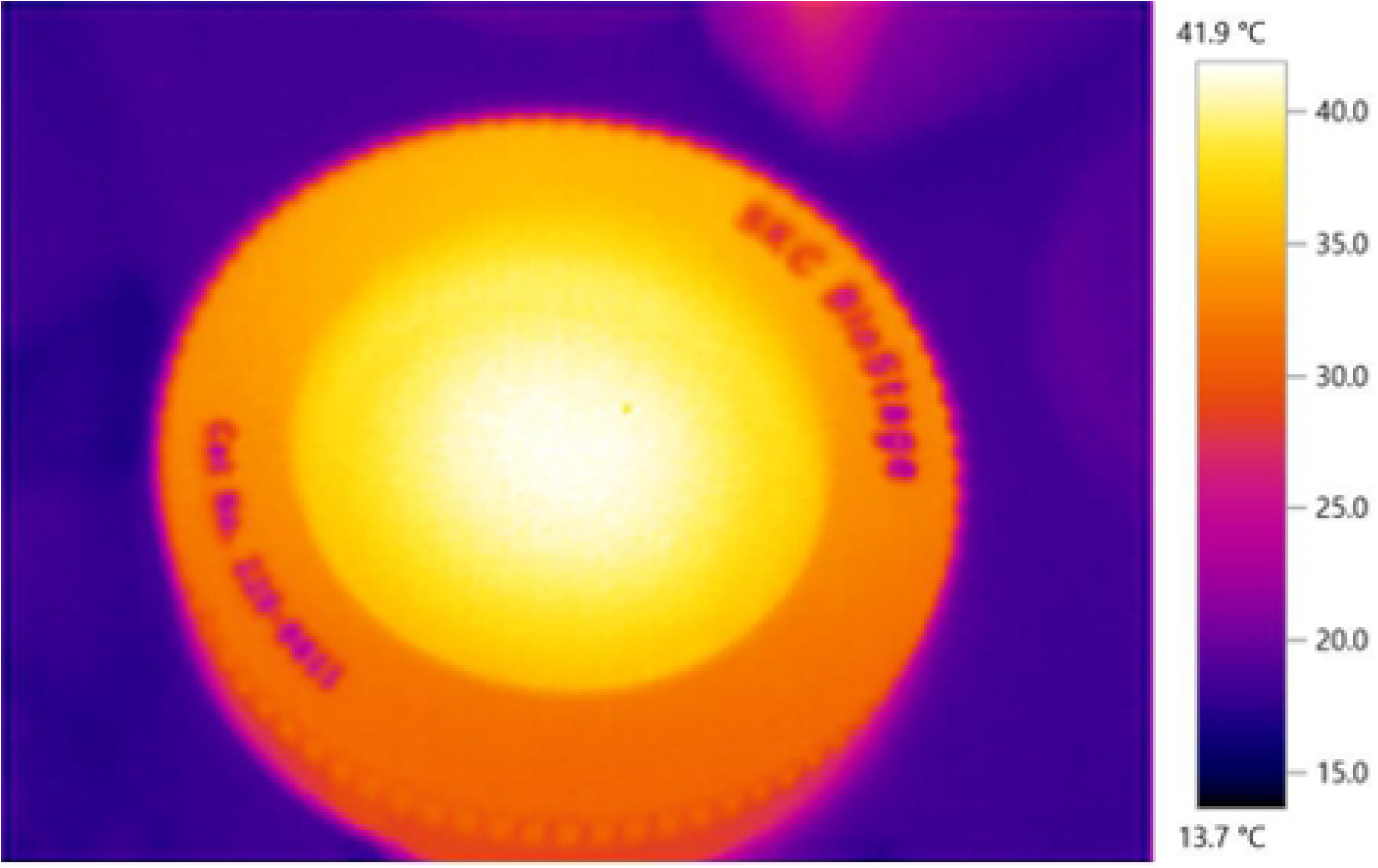
Flame-sanitisation of Andersen 400-hole top-plate. (A) Photograph of results of samples taken by swab from SKC BioStage 400-hole Andersen top-plate doped with an excess of viable dry *Penicillium chrysogenum* spores; (B) Sample after top-plate was liberally doused with common household methylated spirits but not ignited; (C) Sample after top-plate was doused with alcohol and then ignited then allowed to cool for 1 minute before sampling; (D) Thermographs of flaming SKC BioStage 400-hole Andersen top-plate: top-most image was at the time of ignition (maximum estimated temperature 101.9°C); image second from top was 13 s after ignition and continuing to burn; image third from top / at bottom as 35 s after ignition at approximate time when the flames had ceased. It was noted that the temperature of the aluminium top-plate was less than 42°C at this time and was only slightly warm to the touch when repeated on other occasions without added mould spores.

## Discussion

The use of SabCG as commonly formulated and commercially available was found to be reasonably consistent and sensitive for the detection, enumeration and identification of airborne viable fungi including a variety of moulds and yeasts found outdoors, at least in suburban Melbourne, Australia. This was used as a baseline / proxy for the indoor environment, and in the context of a relatively inexpensive, reasonably rapid and difficult to mis-read test. The medium is non-toxic and simple, having an amino-acid/peptide source widely used in microbiology and a glucose/dextrose source, and supports the growth of a wide range of organisms found in outdoor air, which do not require growth factors or such from sources such as malt or vegetable juices, and is not selective for amylase- or maltase-positive organisms, and a wide range of a_w_ requirements, which is useful for its intended purpose in estimating how mouldy a house is, especially those that are damp.

The inclusion of anti-bacterial antibiotics chloramphenicol and gentamycin appears to improve fungal colony morphology, colouration and rate of development by suppressing bacterial growth and hence making enumeration and identification faster and easier even for more experienced staff, including avoiding having to prepare a separate slide for each colony for examination at sufficiently high power magnification to determine if it is a yeast or bacterium.

Detection and enumeration of early-coloniser fungi such as *Penicillium, Aspergillus, Alternaria, Ulocladium, Rhizopus, Mucor* and yeasts may be a useful proxy for the general degree of mouldiness of a house as indicator organisms, being always present in low numbers outdoors (thus a useful control for the sampling equipment, media and culture conditions), growing rapidly and easily, are relatively easily enumerated and identified, are present in significantly elevated numbers in water damaged buildings, and may indeed cause respiratory disease directly.

Further experiments are planned to better analyse statistical aspects of the original 400-hole air sampling method published by Andersen in 1958 [17], as well as more meaningful analyses of real-world houses with and without known mould/moisture issues, and eventually ideally finding any hypothetical correlation between mould and reported symptoms by occupants. This has been especially difficult with the great variety of different media and methods used historically, coupled with a lack of clarity surrounding the different objectives and practical considerations of estimating the mouldiness of an inhabited building, compared with more well-known but significantly different concerns for testing foodstuffs, food-preparation surfaces, surgical and manufacturing clean-rooms, pathology samples including medical, veterinary, plants, etc.

The use of malt-extract-based and/or starch-bearing agar media, often used for the detection of plant pathogens and the contamination of plant-derived foodstuffs, is of questionable suitability for detecting organisms saprophytically degrading organic materials found in damp houses such as carpet, paper, cardboard, plasterboard, timbers, etc., or natural micro-environments such as leaf litter, fallen logs, grass, or animal materials such as hair, wool, fur, skin flakes, leather, dander, etc. This is possibly because such household dusts and materials are unlikely to have significant amounts of starch or maltose compared with foodstuffs and germinated and/or rotting grains.

Maltose does hydrolyse over time and temperature, reportedly approximately 5% or so at 120°C for 1 hour [72], and supported by the results indicating approx. 4.5% after an autoclave cycle of 1 L liquid media. Malt and hence malt-extract is highly variable, being a pivotal aspect of brewing beer and whisky (or whiskey) using different grains including barley, wheat, rye and germinating them under differing conditions to cause the starch to enzymatically break down into various sugars including maltose, and then may be roasted and even smoked to impart a variety of flavours before further processing and extraction typically including concentration by boiling and evaporation, all of which have different, variable and/or un-reported durations and conditions of heat treatment. This would likely lead to significant regional and batch-to-batch variation that is not well controlled or described and may or may not have a significant effect on sampling results when attempting to compare them between groups using different media suppliers, autoclave conditions or working in different countries and/or over time.

Logically and practically it ought to be better to increase consistency of results by reducing batch-to-batch variability by simplifying the media, using chemically-pure glucose (dextrose), and ideally a consistent protein/peptide/amino acid source such as Mycological Peptone™ (Oxoid) or similar, and using reliable stable antibiotics such as chloramphenicol and gentamycin, as per SabCG. The notion that the high a_w_ of the SabCG media might suppress the apparent numbers of outdoor airborne fungi by preventing the growth of numerous xerophilic organisms present, or other possible causes is not supported by the data when the low a_w_ media, DG18 (with antibiotics) was compared with SabCG during winter in Melbourne, Australia.

Other workers had noted a reduction in the viability of some common fungal spores grown under low a_w_ then exposed to high a_w_ media, putatively due to an osmotic-shock effect causing the spores to swell and explode [39,42], thus hypothetically reducing the apparent numbers of airborne fungi recovered on high a_w_ media such as SabCG compared with low a_w_ media such as DG18. This is curious given that few common fungi are markedly inhibited by high a_w_ [73], and because the likely highest contributors to fungal growth are high a_w_ materials/environments, and high a_w_ materials/environments are the notable problem in a damp house and/or WDB. This is presumably quite different to the problems of low a_w_ food spoilage by xerophilic/xerotolerant organisms. Of course, in assessing a building for mould in practice it is to best achieve a reasonable compromise between detecting the full range of viable organisms possibly present, or the subset of ‘indicator organisms’ virtually guaranteed to be present if the building is or has recently been damp and thus mould-affected, and to do so reasonably consistently, rapidly in culture and during enumeration / identification. SabCG appeared to achieve a reasonable balance of this under the experimental conditions and using outdoor airborne fungi as a proxy for the range of organisms found in houses generally or when damp/mouldy.

That the V8c agar medium (with antibiotics) significantly yielded the lowest numbers of outdoor airborne fungi when compared with SabCG and DG18c media in winter in Melbourne, Australia, was interesting but not unexpected given its general paucity of simple sugars and amino acids. That the largest colonies found on the V8c agar medium were always associated with a zone of clearing of the starch granules under and around it was interesting, especially when very small colonies were generally not associated with a zone of clearing. This suggests that the large colonies are able to grow because they were digesting the starch and hence at a considerable advantage in the otherwise relatively energy-poor medium.

The V8c and the DG18c media were hence inferior to the SabCG media for the detection of outdoor airborne fungi, at least in winter in suburban Melbourne, Australia.

The generally-cited incubation conditions for DG18 (25°C, 5-7 days) presented some challenges given that this is often more than the time required for many common moulds to grow to maturity, sporulate and have progeny colonies of a size and state of maturity making them appear to be the originally collected generation, albeit usually smaller but tending to appear like other slower-growing organisms, thus adding a source of bias and confusion. Similarly, other especially sparsely-growing / wispy organisms tend to spread avidly and hence cover smaller colonies, obscuring them and making enumeration and identification difficult. The longer time also presents a problem when there is a potential health-risk at a likely mouldy house, office, etc., and time for results turnaround is important. Hence the use of 3 days incubation at 27°C as standard appeared to be a reasonable compromise, being warm enough to allow the reasonably rapid growth of many organisms, but not so warm as to inhibit temperature-sensitive organisms such as *Penicillium* and *Cladosporium* species commonly found in damp houses. Many environmental organisms including some strains of plant pathogens *Eutypa lata* and *Botryosphaeria* spp. do not have good hyphal growth in culture at temperatures much above 20-24°C depending on the climate they were isolated from [74], but are not the focus of studies of indoor air quality and the determination if a house is mouldy due to water ingress.

In testing various media, it was tempting to use more controlled conditions such as filtered air intentionally seeded with known species of moulds, but it was thought it would be a better test of the natural world to use the likely wider range of organisms found outdoors. Additionally, in testing houses and other buildings for mould, the outdoor air is always tested and compared as a control given that non-mouldy / non-WDB typically have a similar number of airborne as outdoors, but significantly mouldy / WDB have more than outdoors, albeit typically of a narrow range of organisms that grow rapidly in damp conditions on building materials. Therefore, the more important concern is to reliably and rapidly detect the likely relative shift in the range of organisms differentially rather than exhaustively/absolutely.

When the results of the number of outdoor airborne viable fungal CFU/m^3^ detected by culture on standard/stock SabCG medium were compared between each different season (excepting days of low air velocity or an unusually high outlier on 22 January 2020) it was found that the means of the results for spring (Fig 6, September 2019), summer (Fig 8, February 2020), autumn (Fig 2, April 2020) and winter (Fig 4, July 2020) were 282 (SD 22), 316 (SD 30), 289 (SD 19), and 242 (SD 69), respectively. Hence the mean of the means was 282 CFU/m^3^ (SD 30). This is interesting to note as this approximate value is often seen in practice when sampling outdoor air as a control prior to entering a building under assessment for mould, excepting adverse or unusual weather events including rain, strong hot winds, or the air sampling unit being positioned too close to or downwind of a notably mouldy building or materials removed from one, or some types of trees and wetland areas via personal observation of many hundreds of sampling occasions over many years and locations.

While it is possible to flame-sterilise/sanitise the 400-hole Andersen impactor top-plate, it is of questionable advantage to do so while onsite, likely having several orders of magnitude less than 1% an effect on the results even if taken from a very mouldy location to a sterile one, especially compared with the typical natural sampling uncertainty. Flaming onsite carries some practical considerations such as transporting and carrying flammable liquids, and setting fire to it between uses, frequently while wearing flammable gloves and/or disposable polyethylene overalls (e.g., Tyvek™), carrying flammable plastic bags, and often in environments with large amounts of sawdust, cardboard particles, construction materials and waste, plastic sheets used for containment cells during remediation works, paints and thinners, and sometimes quite strong air currents either outdoors, or from outdoors in damaged buildings, or from blowers, air-movers, fans, air-filtering units, heaters, air-conditioning units, dehumidifiers, etc., within buildings often without functional smoke alarms, fire-fighting equipment and functional fire suppression sprinkler systems, proper fire escapes, or often floors, stairs and power.

Previous experience found that collection efficiency was impaired when the top-plate was wiped with cleaning solutions that left any residue such as benzalkonium chloride and detergents, and hence the regular use of hot RO water and an ultrasonic bath was implemented to clean the holes, keeping them open and keeping air flow consistent between uses. It was also found that the number of CFU detected by the 400-hole sampler was not significantly affected by having been used previously in a very mouldy location, such as sampling a ‘clean’ location immediately after a location with high numbers of viable airborne moulds via personal observation of many such sampling occasions. This was not surprising given that the holes are 0.25 mm diameter, and approximately 1.5 mm deep. And thus all 400 holes together have a collective void-volume of less than 30 μL. Hence, for there to be a reasonable chance of increasing a subsequent air sampling run by one single CFU there would have to be one CFU within the 30 μL of void-volume from the previous air sampling run, which would therefore be 1 CFU/30 μL which is 30 × 10^−3^ L, and hence 3.3 × 10^7^ CFU/m^3^, which is very far beyond the typical 282 CFU/m^3^ (343 – 222 at +/- 2SD, hence the 96% confidence interval based on the mean SD) found outdoors normally and even far beyond the lower limit of the highest risk category commonly cited, 5,000 CFU/m^3^ by many orders of magnitude. It was therefore concluded the net effect of sampling even the likely highest possible degree of viable airborne fungal contamination without cleaning the 400-hole top-plate would be significantly less than 1% and hence negligible especially considering the noted greater variation in results from samples taken in the same location using the same media, etc., presumably due to the random nature of airborne mould particles and sampling in general. Prudence and habit, however, meant that the 400-hole top-plate was wiped top and bottom with a commercially available single-use disposable lens cleaning wipe that comes pre-soaked with isopropanol to remove dusts rather than sterilise or sanitise the top-plate. This is reflected in sampling protocols for various airborne microorganisms including fungi and bacteria via the same Andersen sampling apparatus, using isopropanol merely to clean the sampling plate without setting fire to it [75], and ideally the pump unit, hands/gloves and other test equipment potentially exposed to mould, other fungi and organisms in a notably mouldy, dirty or dusty building.

## Acknowledgments

Many thanks are given to all the staff at the Media Preparation Unit, The Peter Doherty Institute for Infection and Immunity, The University of Melbourne, Parkville/Melbourne, Victoria, Australia, with particular acknowledgements (in no particular order) to Elena Paraskeva, Kim Lai Bell, Claire Fraser and Elizabeth Trajcevska, with humble apologies to anyone I have missed.

## Supporting Information

**S1 Tables of raw data**. These tables include CFU/plate counted, calculated CFU/m^3^ air via Andersen, 1958 with a 1.25x adjustment to the raw CFU/plate given polymer Petri dishes were used, and general categories of genera as adapted from ASTM D7391-20 section 12.3.2 [76], pertaining to Figs 1-9.

**S2 Tables, data analysis and graphs**. A MS-Excel notebook of several spreadsheets pertaining to Figs 1-9, including ANOVA analysis, means, standard deviations, and charts/graphs.

